# Spatial Transcriptomics and Single-Nucleus Multi-omics Analysis Revealing the Impact of High Maternal Folic Acid Supplementation on Offspring Brain Development

**DOI:** 10.1101/2024.07.12.603269

**Authors:** Xiguang Xu, Yu Lin, Liduo Yin, Priscila da Silva Serpa, Benjamin Conacher, Christina Pacholac, Francisco Carvallo, Terry Hrubec, Shannon Farris, Kurt Zimmerman, Xiaobin Wang, Hehuang Xie

**Author notes:** These authors contributed equally to this work (Xiguang Xu, Yu Lin).

## Abstract

Folate, an essential vitamin B9, is crucial for diverse biological processes including neurogenesis. Folic acid (FA) supplementation during pregnancy is a standard practice for preventing neural tube defects (NTDs). However, concerns are growing over the potential risks of excessive maternal FA intake. Here, we employed mouse model and spatial transcriptomics and single-nucleus multi-omics approaches to investigate the impact of high maternal FA supplementation during the periconceptional period on offspring brain development. Maternal high FA supplementation affected gene pathways linked to neurogenesis and neuronal axon myelination across multiple brain regions, as well as gene expression alterations related to learning and memory in thalamic and ventricular regions. Single-nucleus multi-omics analysis revealed that maturing excitatory neurons in the dentate gyrus (DG) are particularly vulnerable to high maternal FA intake, leading to aberrant gene expressions and chromatin accessibility in pathways governing ribosomal biogenesis critical for synaptic formation. Our findings provide new insights into specific brain regions, cell types, gene expressions and pathways that can be affected by maternal high FA supplementation.

## INTRODUCTION

Folate, a water-soluble essential vitamin B9, serves as a primary methyl donor in one-carbon metabolism and is involved in various biological processes, including the synthesis and methylation of DNA, RNA and protein molecules [1, 2]. In the brain, folate metabolism facilitates the transfer of one-carbon units for neurotransmitter synthesis and contributes to the regulation of neuronal differentiation and synaptic connectivity [3]. Maternal folate deficiency is known to increase the risk of birth defects, particularly neural tube defects (NTDs) [4]. Consequently, women intending to conceive are advised to supplement with folic acid (FA), the synthetic form of folate, at the dosage of 0.4 mg daily during the periconceptional period [5]. Additionally, the United States and Canada mandated grain fortification with FA since 1998 [6, 7], which has proven effective in reducing the incidence of NTDs at birth in both countries [6, 8]. Due to nationwide grain fortification and the common practice of multivitamin supplementation during pregnancy and lactation, significant increases in folate levels have been observed in maternal serum and breast milk [9, 10]. For instance, following the mandatory grain fortification with FA, serum folate levels in the US population have risen by 2.5 times [9]. Data from the National Health and Nutrition Examination Survey (NHANES) and the Boston Birth Cohort [11] have consistently shown that about a quarter of reproductive age women in the US have circulating folate level in excessive range [12].

There are growing concerns regarding potential adverse impact of excessive maternal FA intake on offspring brain development. Several cohort studies have indicated that elevated maternal FA intake is linked to increased risk of autism spectrum disorder (ASD) [13–15]. Additionally, maternal supplementation with high dose of FA, exceeding the upper limit of 1mg/day, is associated with decreased cognitive development levels in children [16]. In mice, high maternal intake of FA has been shown to cause fetal growth delays [17, 18] and alter cortical neurodevelopment in offspring [19]. Over-supplementation of FA during the periconceptional period has led to abnormal behavioral changes in offspring, including increased anxiety-like behavior, short-term memory deficits, and various forms of learning impairments [18, 20–23]. Furthermore, excess maternal folate supplementation has resulted in morphological changes in offspring brain structures, such as a thinner cortex layer and smaller hippocampus size [18, 19]. Taken together, the growing body of literature regarding excessive FA intake on developing brain underscores the need of further investigating this important public health issue. However, there are significant knowledge gaps. While aberrant changes in gene expression have been noted in the brains of offspring due to excessive maternal folate supplementation [20, 22, 23], we do not know where in the brain, what specific cell types, and which genes/pathways are affected.

Recent advancements in single-cell transcriptomics, such as single-cell and single-nucleus RNA-seq (sc/snRNA-seq), have enabled the generation of transcriptome profiles for specific tissues at single-cell resolution. Given the complex nature of brain tissues, snRNA-seq is particularly advantageous. Despite the absence of cytoplasmic components in isolated nuclei, this method minimizes the risk of aberrant transcription caused by protease digestion or heating [24]. Additionally, coupling snRNA-seq with single-nucleus ATAC-seq (snATAC-seq) allows for the simultaneous detection of gene expression and chromatin accessibility, providing an additional layer of transcriptional regulation at single-cell resolution [25]. However, tissue dissociation in sc/snRNA-seq results in the loss of spatial information [26]. By integrating single-cell and spatial transcriptomics, researchers can capitalize on the strengths of each technology, enabling high-resolution spatial transcriptomic profiling.

In this study, we leverage the unprecedented opportunities afforded by recent advancements in both spatial transcriptomics and single-cell multi-omics analyses to address several crucial questions regarding the impact of maternal high FA intake on offspring brain development. We aim to investigate how excessive maternal FA intake may induce transcriptomic variances across various brain regions, whether various types of brain cells may be affected by high FA differentially, and which gene expressions and pathways are altered and whether they are correlated with changes in chromatin accessibility.

## RESULTS

### FA mouse model and multi-omics sequencing workflow

The FA mice model was adapted from a previous study [19]. Briefly, adult female C57BL/6 mice (8 weeks old) were assigned to receive either a conventional (2 mg/kg, denoted as MF group) or excessive (20 mg/kg, denoted as HF group) folic acid (FA) diet for 2 weeks before mating, continuing throughout pregnancy and lactation. Male pups at postnatal day 21 (P21) were harvested for downstream analysis (**Fig. 1a**). To map gene expression changes across various brain regions, Visium spatial transcriptomics was performed using FFPE coronal sections. Additionally, to discern cell-type specific transcriptional and epigenetic alterations in the hippocampus, single-nucleus multiome analysis was conducted on hippocampal tissues. This approach allows simultaneous profiling of both transcriptome and chromatin accessibility at single-cell resolution (**Fig. 1b**). To evaluate the impact of FA diets on FA levels in mice, we examined the profiles of folate species and one-carbon metabolites in maternal plasma at weaning (P21). The analyzed metabolites included FA, methyltetrahydrofolate (5-Me-THF), homocysteine (Hcy), methionine (MET), cystathionine (CYSTA), S-adenosylmethionine (SAM), and S-adenosyl-L-homocysteine (SAH). Utilizing high-performance liquid chromatography/tandem mass spectrometry (LC-MS), we observed a significant elevated level of FA in the HF group, while no significant difference in the six one-carbon metabolites between the HF and MF groups (**Fig. 1c, Supplementary Fig. 1**). Furthermore, we validated a significant increase in FA levels among both mothers and pups within the HF group compared to those in the MF group using FA ELISA assay (**Fig. 1d&e**).

**Fig. 1.**
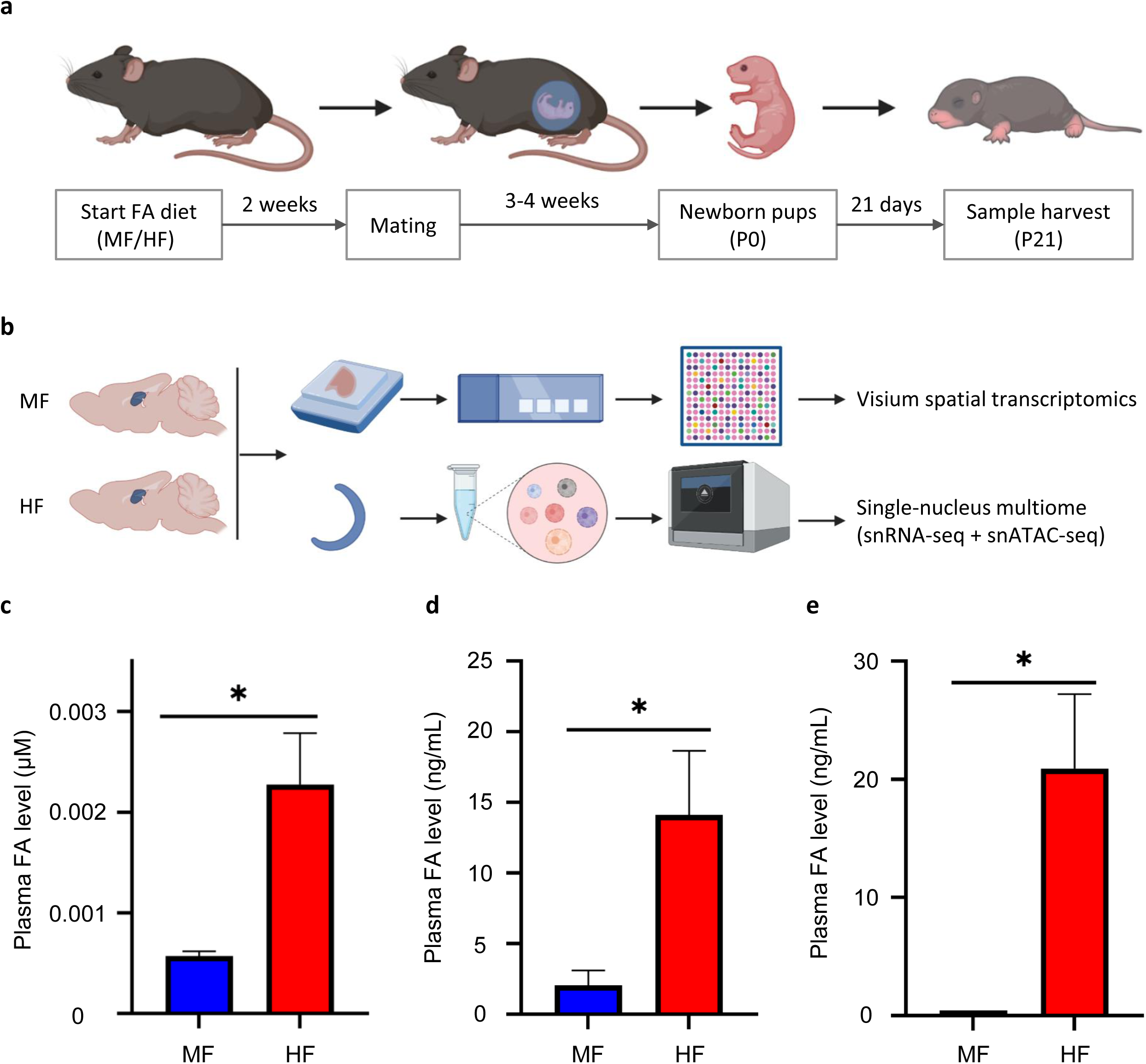
Experimental design. **a**. Adult C57BL/6 female mice were fed with conventional or excessive folic acid (FA) diets (MF/HF) starting two weeks before mating and continuing throughout pregnancy and lactation. Offspring were harvested at P21 for downstream analysis. **b**. FFPE tissue blocks were prepared for Visium spatial transcriptomics. Concurrently, hippocampal tissues were dissected for single-nucleus multiome analysis, generating snRNA-seq and snATAC-seq libraries simultaneously. **c**. Maternal plasma samples were collected when the pups reached P21 and analyzed using LC-MS to quantify folic acid (FA) level. **d,e**. Plasma FA levels were measured by FA ELISA assay in the mother mice (**d**) and P21 pups (**e**). Significant changes are indicated by asterisks (*p<0.05).

### FFPE Visium Spatial transcriptomics recaptures the major brain regions

To capture gene expression profiles across various brain regions, we adopted the Visium spatial transcriptomics approach. For both HF and MF groups, FFPE tissue blocks from two biological replicates were coronally sectioned at 5μm thickness and subjected to Visium spatial transcriptomics analysis using the 10x Genomics Visium-FFPE platform. Each coronal section covered an average of 3,368 spots on the Visium slide, with an average sequencing depth of 158,502,367 million reads, equivalent to a mean of 47,106 reads per spot (**Supplementary Table 1**). To ensure high quality, spots with fewer than 250 unique genes or 500 unique molecular identifiers (UMIs) were excluded from downstream analysis. Consequently, an average of 3,363 spots per sample were retained, with a median of 5,329 genes and 14,184 UMIs per spot (**Supplementary Fig. 2**). To mitigate variations in read coverage among samples, the gene-count matrices were normalized using the counts per million (CPM) method. Clustering the normalized expression matrix using Uniform Manifold Approximation and Projection (UMAP) revealed six major clusters representing the primary anatomical regions of the mouse brain, including the cortex (CTX), hippocampus (HP), thalamus (TH), hypothalamus (HY), striatum (STR), and ventricular system (VS) (**Fig. 2a&b, Supplementary Fig. 4a**). Biological replicates exhibited a robust Pearson’s correlation, with an average r value exceeding 0.9, while the VS regions displayed notable distinctions from other regions (**Supplementary Fig. 3**). We further identified brain region-specific genes (**Fig. 2c, Supplementary Fig. 4b**). For instance, *Satb2* showed high expression in the cortex region, while *Dlk1* exhibited high expression in the hypothalamus region (**Supplementary Fig. 4c**), consistent with previous reports [27, 28]. These findings indicate that the molecular profiling of FFPE mouse brain data accurately reflects the major known anatomical structures. The consistency in expression of region-specific genes observed in our Visium spatial transcriptomics datasets suggests the reliability of spatial transcriptomic analysis conducted in this study.

**Fig. 2.**
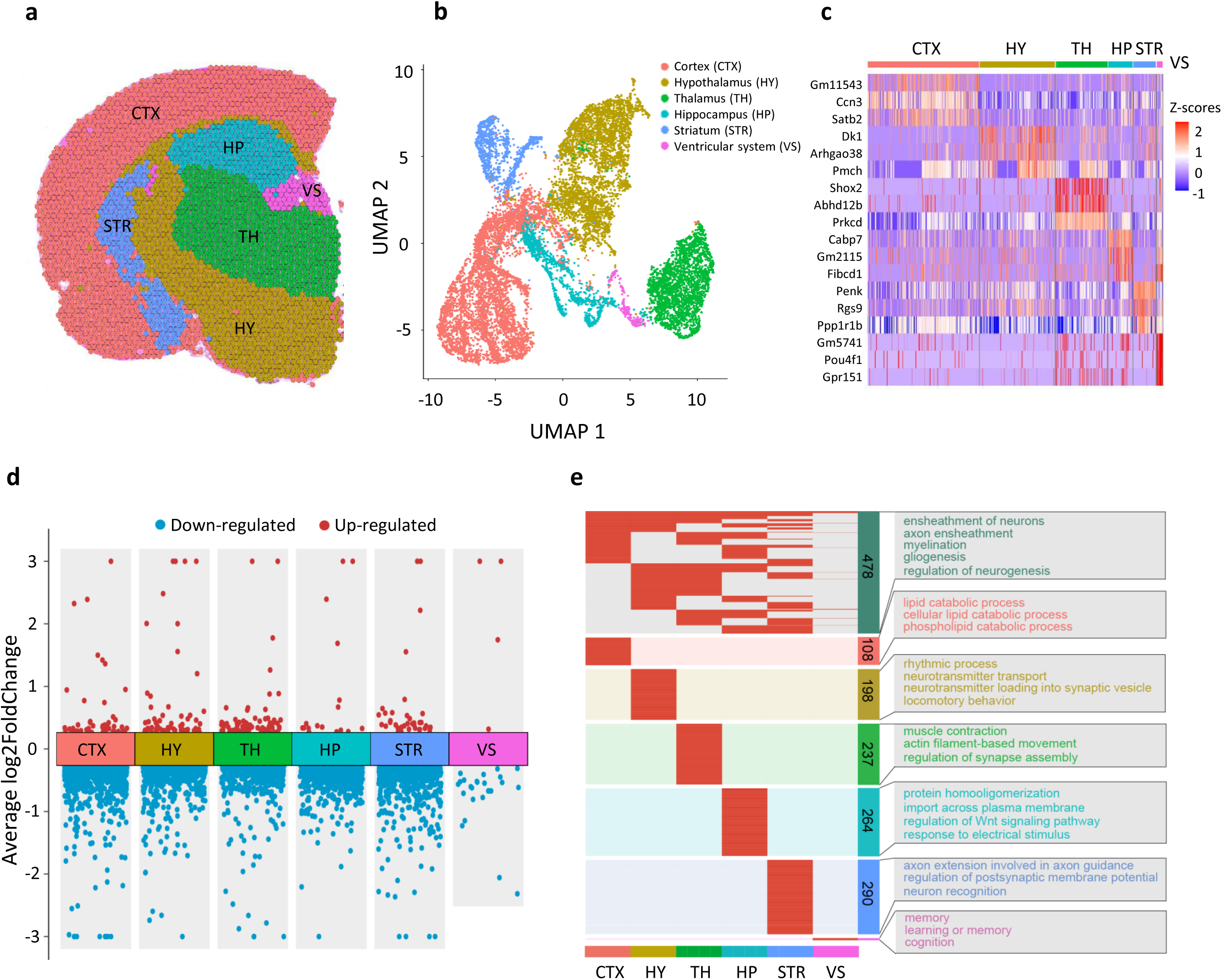
Brain region-specific gene expression changes induced by excess maternal FA supplementation. **a**. Representative feature plot showing the distribution of clusters in the brain sections. **b**. UMAP visualization displaying six major clusters color-coded based on transcriptome profiles. **c**. Heatmap showing the top 3 cluster-enriched genes with Z-transformed average expression levels. **d**. Transformed volcano plot depicting differentially expressed genes between MF and HF for each major cluster. **e.** Gene Ontology (GO) annotation of DEGs identified for each brain region. The DEGs shared by at least 2 brain regions were shown at the top panel.

### Brain region-specific transcriptional changes induced by excess maternal FA supplementation

We next performed brain region-specific differential gene expression analysis. A total of 1,583 differentially expressed genes (DEGs) emerged from the comparison between the HF and MF groups, consisting of 160 upregulated genes and 1,425 downregulated genes (**Fig. 2d, Supplementary Table 2**). We examined the intersection of the DEGs across the six primary brain regions and observed 69.4% of the DEGs showed specificity to certain brain regions (**Supplementary Fig. 5**). Among all the DEGs, 478 genes were common in at least two of the six brain regions, while 1,105 were specific to particular regions. For instance, there are 108 DEGs specific to the cortex, 264 DEGs specific to the hippocampus, and 290 DEGs specific to the striatum (**Supplementary Fig. 5**). This finding suggests that excess maternal FA supplementation induces brain region-specific changes gene expression in the offspring.

To further investigate the biological functions associated with the identified DEGs, we utilized the clusterprofiler package for GO annotation [29]. Functional enrichment analysis revealed a widespread influence of maternal FA intake. For instance, processes such as ensheathment of neurons, axon ensheathment, myelination, gliogenesis and regulation of neurogenesis exhibited enrichment in DEGs shared by multiple brain regions (**Fig. 2e**), highlighting the crucial role of folate in neuronal axon myelination. This finding aligns with previous research indicating folate’s influence on oligodendrocyte survival, axon regeneration [30, 31] and neurogenesis regulation [19, 32]. Notably, DEGs specially identified in the hypothalamus showed enrichment in processes related to neurotransmitter transport and neurotransmitter loading into synaptic vesicle, DEGs uniquely identified in the thalamus are associated with regulation of synapse assembly, DEGs exclusively identified in the ventricular system enriched in processes related to learning and memory (**Fig. 2e**), suggesting that region-specific effects induced by excessive maternal FA supplementation could elucidate the multifaceted consequences of such intake on offspring behavior, among other outcomes. In summary, the aforementioned findings delineate a spatial pattern of extensive transcriptional alterations across various brain regions triggered by excessive maternal FA supplementation.

### Cortex and hippocampus exhibit subregion-specific transcriptomic changes by excess maternal FA supplementation

We then focused our analysis on two crucial brain regions: the cortex and hippocampus, renowned for their essential roles in cognitive functions [26]. Four distinct sub-clusters were identified within the cortex, labeled as Layer 1 (L1), layer 2-3 (L2-3), layer 4-6 (L4-6), and olfactory areas (OLF) (**Fig. 3a&b**). Similarly, sub-clustering of the hippocampus revealed an additional four sub-clusters: CA1, CA2-3, Dentate Gyrus (DG), and a matrix encompassing the stratum oriens, stratum radiatum, and stratum lacunosum-moleculare (**Fig. 3d&e**). Specific markers corresponding to distinct layers were identified in cortex sub-regions. For instance, *Fezf2* exhibited high expression levels in the deep layer 4-6, while *Stard8* showed prominent expression in the outer layer 2-3 (**Fig. 3c, Supplementary Fig. 6a&c**), consistent with previous reports [33, 34]. Similarly, subregion-specific markers were identified in the hippocampus subregions. Notably, established hippocampal subregion markers, such as Spink8, Ptgs2, and Dsp, demonstrated high expression levels in the CA1, CA2-3, and DG regions, respectively [35, 36] (**Fig. 3f, Supplementary Fig. 6b&d**).

**Fig. 3.**
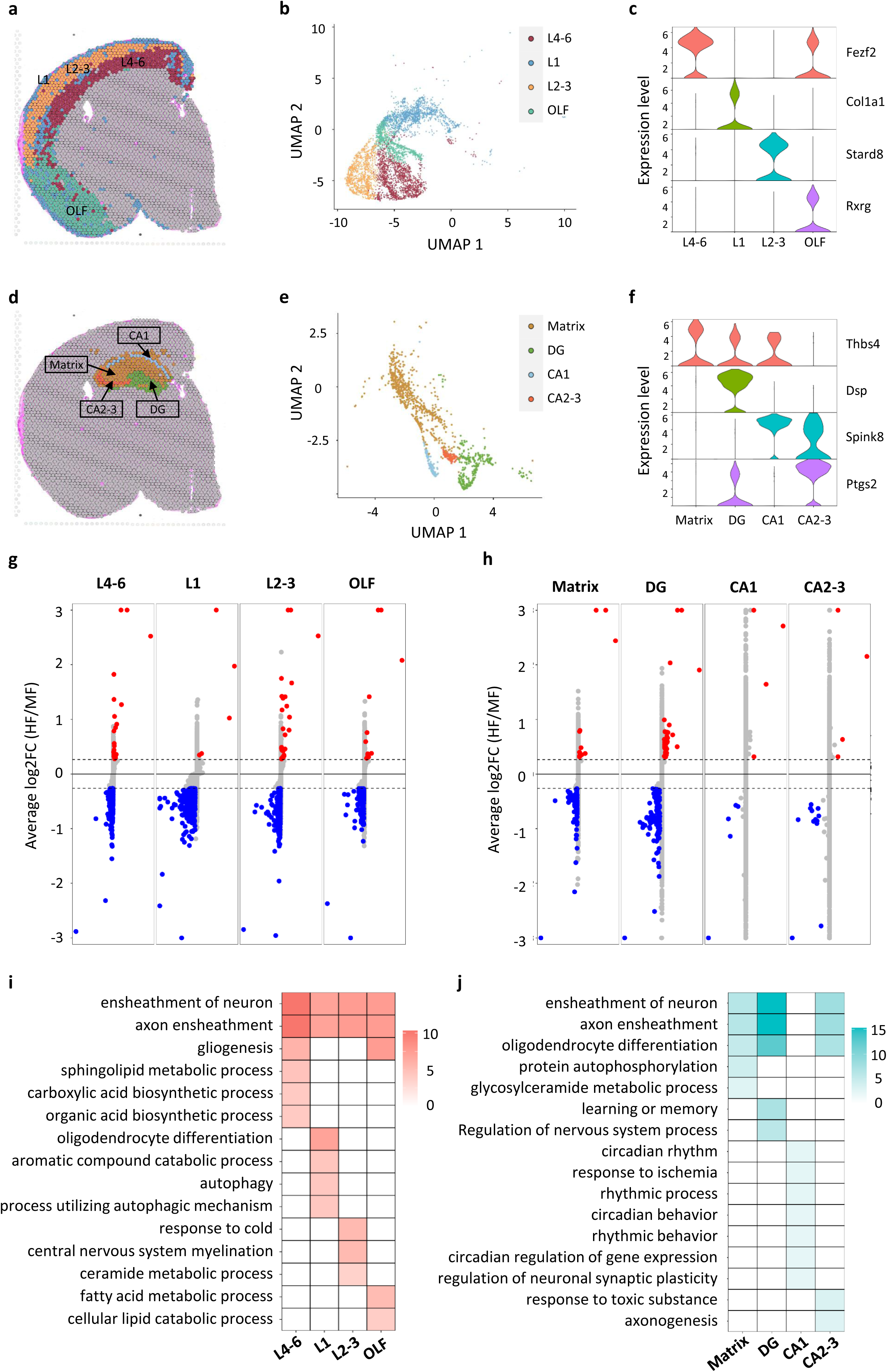
Subregion-specific gene expression changes in cortex and hippocampus induced by excess maternal FA supplementation. **a**. Representative feature plot showing sub-clusters in the cortex region. **b**. UMAP plot categorizing dots into four sub-clusters based on gene expression profiles. L1 (layer 1), L2-3 (layer 2-3), L4-6 (layer 4-6), OLF (olfactory region). **c**. Violin plot showing the expression of marker genes for each sub-cluster in the cortex. **d**. Representative feature plot showing sub-clusters in the hippocampus region. **e**. UMAP plot classifying dots into four sub-clusters based on gene expression profiles in the hippocampus: CA1, CA2-3, DG (dentate gyrus), and matrix. **f**. Violin plot showing the expression of marker genes for each sub-cluster in the hippocampus. **g.** Transformed volcano plot depicting differentially expressed genes (DEGs) between the MF and HF groups for each sub-cluster in the hippocampus. **h**. Transformed volcano plot illustrating DEGs between the MF and HF groups for each sub-cluster in the cortex. **i**. GO annotation of DEGs for each sub-cluster in the cortex region. **j**. GO annotation of DEGs for each sub-cluster in the hippocampus region.

For each sub-cluster within the cortex and hippocampus, we conducted a differential gene expression analysis to elucidate the transcriptional alterations within these subregions. In the cortex, we observed 18 up-regulated genes and 597 down-regulated genes in the L4-6 sub-cluster, 3 up-regulated genes and 593 down-regulated genes in the L1 sub-cluster, 25 up-regulated genes and 576 down-regulated genes in the L2-3 sub-cluster, and 3 up-regulated genes and 198 down-regulated genes in the OLF sub-cluster (**Fig. 3g, Supplementary Table 3**). This highlights a predominance of down-regulated DEGs, suggesting a prevalent suppression of gene expression in the cortex region due to excessive FA supplementation. In the hippocampus, we identified 10 up- regulated genes and 68 down-regulated genes in the L4-6 sub-cluster, 36 up-regulated genes and 103 down-regulated genes in the DG sub-cluster, 3 up-regulated genes and 5 down-regulated genes in the CA1 sub-cluster, and 3 up-regulated genes and 110 down-regulated genes in the CA2-3 sub-cluster (**Fig. 3h, Supplementary Table 3**). Consistently, we observed a similar trend with a higher number of down-regulated DEGs. Notably, the DG sub-region appears to be particularly impacted by excessive maternal FA supplementation in the hippocampus. Neuronal axon ensheathment consistently emerged as the top enriched GO term across all subregions of the cortex and hippocampus, except for the CA1 subregion (**Fig. 3i&j**), probably due to the low number of DEGs identified in CA1. In addition to the shared GO terms among different subregions, we also identified subregion-specific GO terms. For instance, organic acid biosynthetic process was enriched in L4-6, oligodendrocyte differentiation in L1, ceramide metabolic process in L2-3, and fatty acid metabolic process in OLF. Similarly, in the hippocampus, protein autophosphorylation was enriched in the matrix, learning or memory in the DG, regulation of neuronal synaptic plasticity in the CA1, and axongenesis in the CA2-3. Taken together, these findings highlight both shared and subregion-specific effects induced by excessive maternal FA intake.

### Single-nucleus multiome analysis of the P21 offspring’s hippocampus

To further decipher the cell-type specific regulation of gene expression influenced by excess maternal FA intake, we performed single-nucleus multiome analysis on the hippocampus of P21 male offspring from mothers exposed to either MF or HF diets. Our multiome libraries met quality standards and demonstrated a clear separation between cells and non-cells (**Supplementary Fig. 7a**). ATAC peak enrichment was centered at transcription start site (TSS) (**Supplementary Fig. 7b**), with fragment sizes peaking at 150bp (**Supplementary Fig. 7c**). We observed excellent reproducibility across both biological replicates in snRNA-seq and snATAC-seq analyses (**Supplementary Fig. 8a-d**). After filtering out low-quality nuclei, we obtained a total of 11,148 single-nucleus multiome profiles (snRNA-seq and snATAC-seq) from the four hippocampal samples. In our snRNA-seq datasets, we observed a median of 7,260 counts and 2,931 genes per nucleus, while in snATAC-seq datasets, we detected a median of 10,945 counts and a TSS enrichment score of 6.41 (**Supplementary Fig. 8e-h**).

Dimension reduction was performed on each snRNA-seq and snATAC-seq dataset. We applied UMAP and unsupervised clustering using the Approximate Nearest Neighbor (ANN) method to integrate snRNA-seq and snATAC-seq datasets [37], resulting in a total of 13 distinct cell clusters (**Fig. 4a, Supplementary Fig. 9**). RNA expression levels and gene activity scores of marker genes were identified for each major cell type based on snRNA-seq and snATAC-seq results, respectively (**Fig. 4b&c**). We identified six major hippocampal cell types: excitatory neurons (EXC, 6,333 nuclei), inhibitory neurons (INH, 929 nuclei), astrocytes (AST, 1,401 nuclei), oligodendrocyte progenitor cells (OPC, 9937 nuclei), oligodendrocyte (ODC, 869 nuclei), and microglia (MG, 679 nuclei). The identity of each cell type was confirmed using known cell-type specific marker genes: *Slc17a7* for EXC, *Gad2* for INH, *Aqp4* for AST, *Cspg4* for OPC, *Mog* for ODC, and *Ptprc* for MG (**Fig. 4d&e**). Robust correlations were observed between Dimension reduction was performed on each snRNA-seq and snATAC-seq dataset. We applied UMAP and unsupervised clustering using the Approximate Nearest Neighbor (ANN) method to integrate snRNA-seq and snATAC-seq datasets [35], resulting in a total of 13 distinct cell clusters (Fig. 4a, Fig. S8). RNA expression levels and gene activity scores of marker genes were identified for each major cell type based on snRNA-seq and snATAC-seq results, respectively (Fig. 4b&c). We identified six major hippocampal cell types: excitatory neurons (EXC, 6,333 nuclei), inhibitory neurons (INH, 929 nuclei), astrocytes (AST, 1,401 nuclei), oligodendrocyte progenitor cells (OPC, 9937 nuclei), oligodendrocyte (ODC, 869 nuclei), and microglia (MG, 679 nuclei). The identity of each cell type was confirmed using known cell-type specific marker genes: Slc17a7 for EXC, Gad2 for INH, Aqp4 for AST, Cspg4 for OPC, Mog for ODC, and *Ptprc* for MG (Fig. 4d&e). Robust correlations were observed between gene accessibility and gene expression in both the MF and HF groups, with Pearson’s r values of 0.73 and 0.74, respectively (**Fig. 4f&g**). Additionally, correlations across various cell types were explored, revealing the lowest correlation of around 0.57 in astrocytes and microglia, while excitatory neurons exhibited the highest correlation at 0.71 (**Supplementary Fig. 10**). The positive correlation between chromatin accessibility and gene expression underscores that chromatin accessibility is a reliable marker for the regulation of gene transcription.

**Fig. 4.**
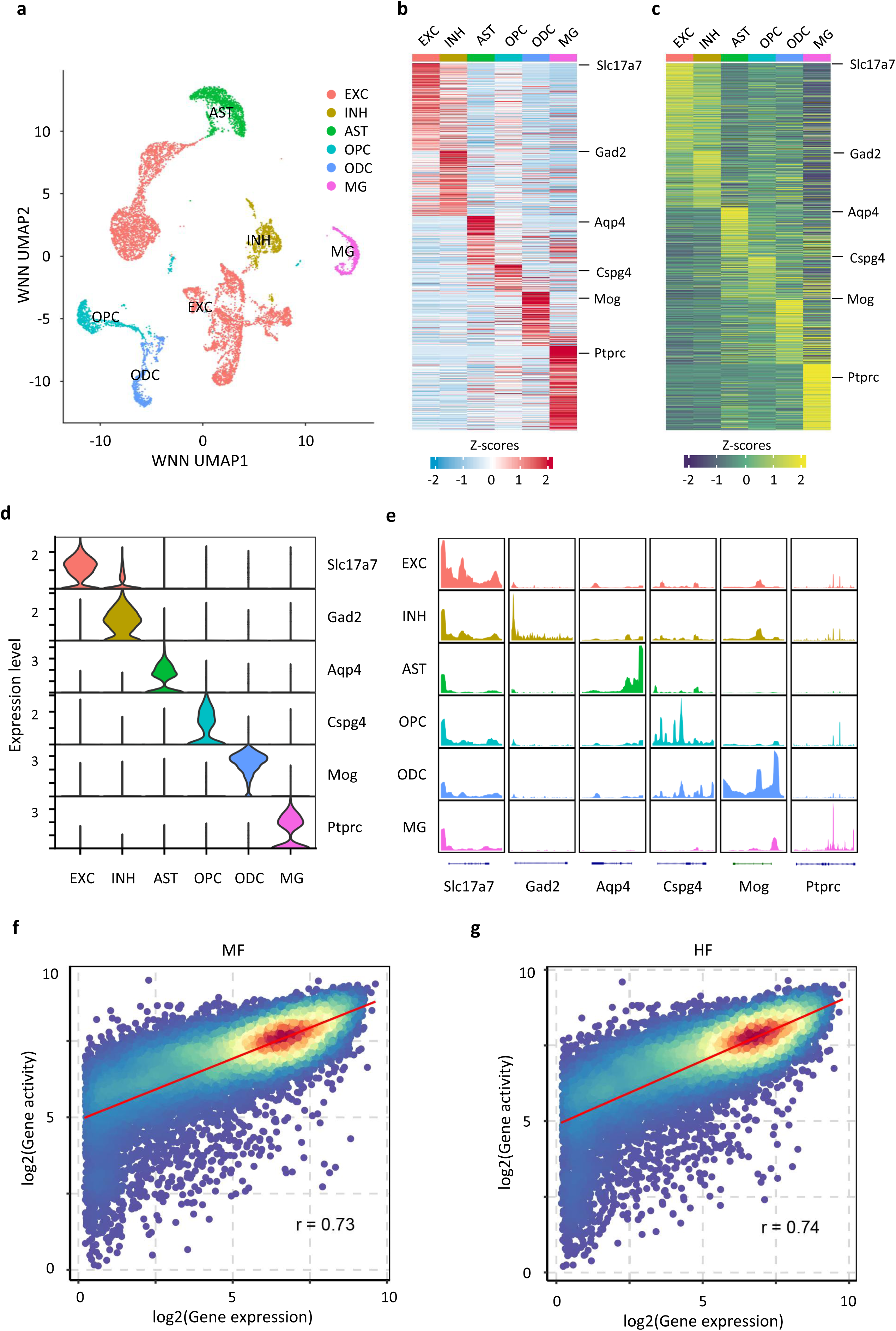
Identification of diverse hippocampal cell types at P21 using single-nucleus multiome analysis. **a**. Integrated UMAP visualization of major hippocampal cell types from 10,234 nuclei profiled for both transcriptome and chromatin accessibility using single-nucleus multiome data. Identified cell types include excitatory neurons (EXC), inhibitory neurons (INH), astrocytes (AST), oligodendrocyte progenitor cells (OPC), oligodendrocytes (ODC), and microglia (MG). **b**. Row-normalized heatmaps of single-nucleus gene expression for cell-type specific marker genes. **c**. Row-normalized heatmap of gene activity for cell-type specific marker genes. **d**. Violin plot showing gene expression levels of marker genes for each major cell type. **e**. Psedo-bulk chromatin accessibility profiles at marker genes for each major cell type. **f, g**. Correlation between gene expression and gene accessibility for MF (**f**) and HF (**g**). Each dot represents a gene, with colors indicating dot density; red indicates high density, and blue indicates low density.

### Excess maternal FA supplementation induced cell-type specific changes in gene expression and chromatin accessibility

Excessive FA intake during pregnancy and lactation has been shown to cause aberrant gene expression changes in the brain tissue of offspring [3]. However, the understanding of cell-type-specific gene expression changes remains limited. To address this gap, we conducted comparative transcriptome and epigenome analyses between the MF and HF groups using single-nucleus multiome datasets derived from hippocampal tissues. First, we examined whether excessive maternal FA intake leads to shifts in the population of major cell types within the hippocampus. Our analysis of the proportions of the six major cell types between the MF and HF groups revealed no significant changes (**Supplementary Fig. 11**). We next conducted differential gene expression analysis between the MF and HF groups within the six major cell types, identifying a total of 173 DEGs. Among these, 9 DEGs were detected in three or more cell types, suggesting that excessive maternal FA intake can influence multiple cell types (**Fig. 5a, Supplementary Table 4**). However, a notable proportion of DEGs, 156 in total, were specific to excitatory neurons. Interestingly, the expression of these DEGs displayed a moderate correlation with the gene activity score, as indicated by ATAC peaks in promoter and gene body regions (**Fig. 5b**). Subsequently, we focused on DEGs identified within excitatory neurons and performed gene ontology (GO) analysis. It revealed a significant enrichment of genes involved in translation, ribosomal small and large unit biogenesis, and neuronal synaptic plasticity (**Fig. 5c**). For example, *Cmss1* and *Filip1l* are two genes related to ribosomal biogenesis and cell proliferation, respectively [38, 39]. Although the median expression of *Filip1l* in excitatory neurons did not differ between HF and MF groups (**Fig. 5d**), lower expression level of these two genes were observed at the 90^th^ percentile among all cells in the HF group compared to the MF group (**Fig. 5e**). Additionally, a substantial difference in the pseudo-bulk ATAC-seq signal was observed in the genic regions of these two genes between the two groups of excitatory neurons (**Fig. 5f**). This suggests that epigenetic mechanisms may serve as intermediate regulators in response to folate, subsequently influencing gene expression changes.

**Fig. 5.**
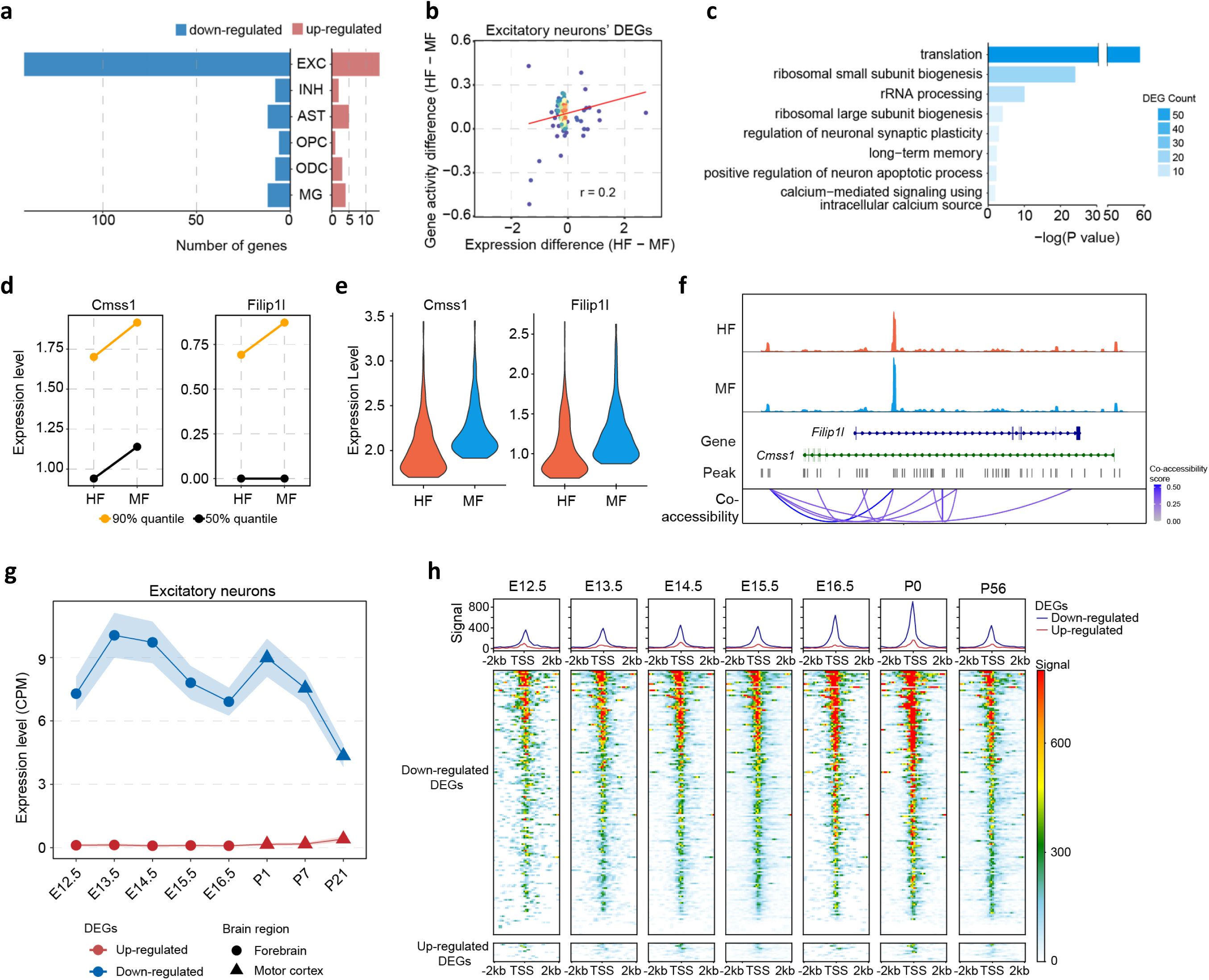
Cell-type specific transcriptional changes induced by excess maternal FA intake. **a**. Number of differentially expressed genes (DEGs) identified between HF and MF group for each cell type. **b**. Correlation between gene expression change and gene activity change between HF and MF group for excitatory neurons. **c**. Functional enrichment analysis for DEGs in excitatory neurons. **d**. Expression of Cmss1 and Filip1l in excitatory neurons of HF and MF. 50% quantile of expression values are colored in black and 90% quantile are colored in orange. **e**. Violin plot shows the expression level of Cmss1 and Filip1l in top 10% expressed excitatory neurons of HF and MF, separately. **f**. ATAC-seq signal surrounding Cmss1 and Filip1l genes in HF and MF. **g**. Expression of excitatory neuron DEGs during brain development. **h**. ATAC-seq signal in the promoters of DEGs along with brain development.

To explore the expression change and chromatin accessibility dynamics of DEGs in excitatory neurons during brain development, we utilized publicly available single-cell RNA-seq [40, 41] and single-nucleus ATAC-seq [42] datasets from multiple embryonic and postnatal brain developmental stages. We extracted the expression data of excitatory neurons from these datasets to analyze the developmental changes in gene expression and chromatin accessibility for these DEGs. Our analysis revealed that the up-regulated DEGs maintained a stable, low level in both expression and gene accessibility (**Fig. 5g&h**). More interestingly, a sharp decline in both expression and gene accessibility was observed for the downregulated DEGs during postnatal stages, indicating the functional importance of these DEGs in embryonic neuronal development compared to postnatal stages.

### Multi-omics integration reveals spatial distribution of hippocampal excitatory neuron subtypes influenced by maternal FA intake

To investigate the effects of excess maternal FA intake on excitatory neurons in the hippocampus, we re-clustered the 6,333 Slc17a7+ excitatory neurons using the top 2,000 highly variable genes. This analysis unveiled five distinct excitatory neuron subtypes: CA1 neurons, CA3 neurons, DG mature neurons, DG immature neurons, and Erbb4+ neurons (**Fig. 6a**). We identified RNA expression levels and gene activity scores for marker genes specific to each excitatory neuron subtype (**Supplementary Fig. 12a&b**). Representative marker genes for each subtype were illustrated in both RNA expression levels and gene activity scores (**Fig. 6b&c, Supplementary Fig. 12c**). Cellular composition analysis indicated no significant alterations in the proportion of these excitatory neuron subtypes (**Supplementary Fig. 13**).

**Fig 6.**
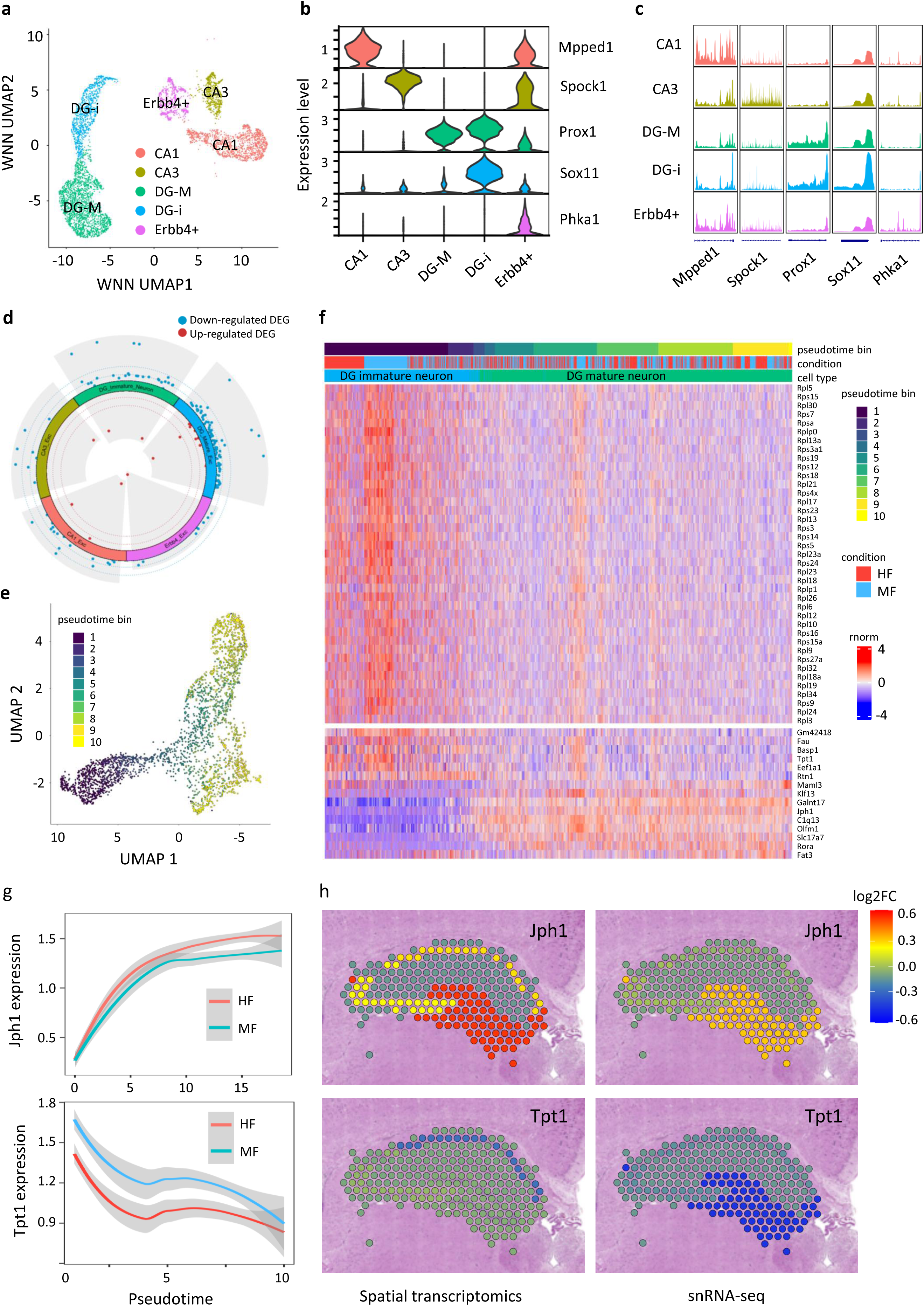
Transcriptional changes in excitatory neuron subtypes induced by excess maternal FA intake. **a**. Integrated UMAP visualization of 6,333 hippocampal excitatory neurons profiled for both transcriptome and chromatin accessibility using snRNA-seq and snATAC-seq data. Hippocampal excitatory neuron subtypes include CA1 neurons (CA1), CA3 neurons (CA3), DG immature neurons (DG-i), DG mature neurons (DG-M), Erbb4+ neurons (Erbb4+). **b**. Violin plot illustrating gene expression levels of marker genes for each hippocampal excitatory neuron subtype. **c**. Pseudo-bulk chromatin accessibility profiles at marker genes for each hippocampal excitatory neuron subtype. **d**. Differentially expressed genes (DEGs) identified between the MF and HF groups for each hippocampal excitatory neuron subtype. **e**. UMAP showing the trajectory analysis of DG immature and mature neurons. **f**. Heatmap showing the expression profiles of trajectory-related DEGs along the trajectory of DG neurons. **g**. Expression of Jph1 and Tpt1 in MF and HF groups along the trajectory of DG neurons. **h**. Spatial profiles of Jph1 and Tpt1 derived from spatial transcriptomics datasets or snRNA-seq datasets projection to spatial transcriptomics, represented by log2 fold change of HF/MF.

We further conducted differential gene expression analysis within the five excitatory neuron subtypes. Of the 109 DEGs identified between the MF and HF groups, 84 (77.1%) overlapped with DEGs found in excitatory neurons across the entire hippocampal region. Additionally, DEGs identified in various hippocampal subregions were enriched in the DG region (**Fig. 6d, Supplementary Table 5**), which is known for its role in neurogenesis, actively generating new neurons [43]. Both immature and mature excitatory neurons in the DG region exhibited significant alterations in these DEGs, suggesting that DG neurons are more vulnerable to aberrant maternal FA intake. It’s noteworthy that a substantial portion of the DEGs (20 out of 29 in DG immature neurons and 32 out of 90 in DG mature neurons) were ribosomal protein-coding genes involved in translation, indicating a critical influence of folate on RNA translation in DG neurons.

Given that DG neurons can be categorized into immature and mature groups, we combined the two groups for trajectory analysis (**Fig. 6e, Supplementary Fig. 14a**). Surprisingly, seventy-eight ribosomal protein-coding genes were identified as pseudotime-related genes, underscoring the importance of ribosomal biogenesis in the maturation process of DG neurons. We also investigated how the DEGs identified between DG immature and mature neurons overlap with genes related to pseudotime, revealing a shared set of 39 ribosomal protein-coding genes. These differentially expressed ribosomal protein coding genes showed highest expression levels in DG immature neurons (**Supplementary Fig. 14b**). This expression decreased as neurons in the DG region matured, with the lowest levels observed in other hippocampal excitatory neuron subtypes (**Supplementary Fig. 14b**). Along the trajectory, ribosomal protein-coding genes exhibited a similar dynamic expression pattern, with a decrease in expression as maturation approached (**Fig. 6f**). To validate the expression patterns of these ribosomal genes during neuronal development, we utilized aforementioned publicly available single-cell RNAseq data from forebrain and motor cortex regions, extracting and aggregating expression data specific to excitatory neurons [40, 41]. The expression of these ribosomal genes exhibited highly dynamic profiles, with an overall decreasing trend throughout neuronal maturation, particularly during the postnatal stages from P7 to P21 (**Supplementary Fig. 14c**). These results suggest excessive maternal FA intake results in aberrant expression of ribosomal protein-coding genes, potentially impacting RNA translation.

Lastly, we investigated whether the alterations observed at the single-nucleus level could be detected in the spatial organization of the brain. The 10X Visium platform, which features spot sizes of 55 μm covering 1-10 cells, derives expression profiles from a mixture of both nuclear and cytoplasmic mRNA populations. In contrast, snRNA-seq data offers single-cell resolution but is limited to nuclear mRNAs. Despite these differences in RNA sources and cellular resolution, we correlated the snRNA-seq data with Visium spatial transcriptomics data using marker genes. This correlation revealed the clear spatial distribution of hippocampal excitatory neuron subtypes (**Supplementary Fig. 15**). We then projected the DEGs identified in the snRNA-seq data onto the spatial map, focusing on the differentially up-regulated and down-regulated genes in DG neurons between the HF and MF groups. For instance, *Jph1* was up-regulated while *Tpt1* was downregulated in the HF group (**Fig. 6g**). Consistent patterns of expression changes for these DEGs were observed in both the spatial transcriptomics and snRNA-seq datasets (**Fig. 6h**). The comparatively subdued changes in the spatial transcriptomics data likely result from signal dilution due to the presence of other cell types. Nevertheless, the congruent spatial expression patterns observed between the spatial transcriptomics and single-nucleus transcriptome data suggest successful integration, particularly for DEGs highly expressed in specific cell subtypes.

## DISCUSSION

In the post-fortification era, population-wide plasma folate levels have significantly increased [9]. Accumulating evidence suggests a potential risk associated with excessive FA intake, particularly during the periconceptional period [14, 19, 22, 44]. In this study, we have generated a reference dataset that offer valuable insights into the spatial and single-cell resolution of transcriptional changes in the brains of offspring resulting from excessive maternal FA supplementation. By utilizing multi-omics data from various levels, we have uncovered several noteworthy findings. Firstly, excessive maternal FA intake broadly impacts multiple brain regions, affecting gene pathways involved in neurogenesis, neurotransmitter transport, and neuronal axon myelination. Specifically, myelination associated terms were shared across multiple brain regions, this is in line with previous finding that folate metabolism regulates oligodendrocyte myelination during central nervous system development [30]. Additionally, excessive maternal FA supplementation also induces brain region-specific effects on the expression of genes involved in the learning and memory processes. This suggests that different types of brain cells or cellular communities may respond differently to FA intake, with these differences potentially arising as direct or indirect consequences of suboptimal FA intake during brain development.

Secondly, high maternal FA intake induced expression changes in ribosomal protein-coding genes, including both large subunits and small subunits. During protein synthesis, the small subunit of a ribosome identifies codons by binding to the mRNA template and choosing aminoacyl-tRNAs matching the mRNA codons, while the large ribosomal subunit facilitates the crucial chemical step of peptide bond formation [45]. This suggests a targeted influence on the translation of particular gene subsets alongside broader changes in protein synthesis. Interestingly, previous studies revealed a regulatory role of folate metabolism in RNA translation [46, 47]. Throughout brain development, ribosomal proteins are abundant, yet their abundance diminishes as the brain reaches maturity. Such a reduction in neuronal ribosomal proteins aligns with the decrease in the inherent ability of neuronal axons to grow as they reach their post-synaptic targets and establish operational synaptic connections [48]. The contrasting correlation between the maturation status of cells and the expression levels of ribosomal proteins has been previously documented in normal hematopoietic cells [49]. During postnatal development, synapse genesis is tightly regulated by a combination of genetic programs, neural activity, and environmental factors. Disruptions to this process can have profound effects on brain function and may contribute to neurodevelopmental disorders. Importantly, at P21, the mouse brain is in a dynamic state of development, with synapse formation playing a pivotal role in shaping neural circuits and networks. This period is characterized by a high level of synaptic plasticity, where synapses are both forming and refining in response to various environmental and developmental cues. Thus, our findings imply that excessive maternal FA intake may modify the expression of ribosomal protein genes, consequently interrupting the neuronal maturation process and impairing synaptic genesis.

Lastly, our single-cell multi-omics analysis revealed a robust correlation, with Pearson’s r values of 0.74, between gene expression and chromatin accessibility across all genes. However, this correlation substantially diminishes when assessing each gene across various cells. For DEGs, the difference in gene expression and the difference in chromatin accessibility shows a moderate correlation with Pearson’s r values of 0.2. This suggests that while chromatin accessibility plays a pivotal role in regulating gene expression, mechanisms other than chromatin accessibility likely participate in gene expression regulation. Further research is warranted to elucidate alternative mechanisms by which one-carbon metabolism may influence gene expression.

## MATERIALS AND METHODS

### Animal husbandry and FA diets

Animal experiments were conducted with the prior approval from the Institutional Animal Care and Use Committee (IACUC) of Virginia Tech. C57BL/6J mice, obtained from Jackson Laboratory, were housed in a standard pathogen-free environment with a 12-hour light/dark cycle. Adult female mice (8-12 weeks old) were randomly assigned to either a control folic acid (FA) diet (2mg/kg) or an excessive FA diet (20mg/kg) for two weeks before mating with males. Clifford/Koury-based L-amino acid defined rodent diets containing either control FA (2 mg/kg) or excess FA (20 mg/kg) without succinyl sulfathiazole were purchased from Dyets Inc. (Bethlehem, PA). These diets were maintained throughout pregnancy and lactation. Pups were harvested on postnatal day 21 (P21).

### Quantification of folate metabolites by Mass Spectrometry

Mice were anesthetized with isoflurane inhalation. Blood samples were collected and plasma was then separated from blood cells by centrifugation at 2,000g for 10min at 4°C using a plasma separator tube (fisher, cat# 02-675-187). An aliquot of 100 μL plasma samples were used for high-performance liquid chromatography/tandem mass spectrometry (LC-MS) to measure the levels of folate metabolites: folic acid (FA), methyltetrahydrofolate (5-Me-THF), homocysteine (Hcy), methionine (MET), cystathionine (CYSTA), S-adenosylmethionine (SAM), and S-adenosyl-L-homocysteine (SAH). Briefly, A mixed standard solution of the 7 metabolites was prepared with their standard substances in water, at 50 μM for each compound. This standard solution was further diluted to have 9 calibration solutions. 20 μL of each plasma or each calibration solution was mixed with 20 μL of an internal standard (IS) solution. The mixtures were vortexed for 1 min and then sonicated in an ice-water bath for 30 s. The solutions were centrifuged at 21,000 g and 5 °C for 10 min. 10-μL aliquots were injected to run LC-MRM/MS on an Agilent 1290 UHPLC system coupled to an Agilent 6495C QQQ mass spectrometer which was equipped with an electrospray ionization (ESI) source and operated in the positive ion mode. Linear-regression calibration curves of the analytes were constructed with the data acquired from the calibration solutions.

### FA ELISA Assay

FA levels in plasma samples were quantified using the FA ELISA Assay kit (Cell Biolabs, cat# MET-5068). Initially, the plate was coated with folic acid conjugate at 4°C overnight. The following morning, the plate was washed with 1x PBS and blocked with assay diluent for 1h at room temperature (RT). Plasma samples or folic acid standards were added to the corresponding wells and incubated at RT for 10min, followed by the addition of anti-folic acid antibody for a 1- hour incubation at RT. After three washes with 1x Wash Buffer, the secondary antibody-HRP enzyme conjugate was added and incubated at RT for 1h. Following three additional washes, the substrate solution was added and incubated for 10min. The reaction was stopped by adding the Stop Solution, and absorbance was measured at a wavelength of 450nm using the Cytation 5 microplate reader (BioTek, serial#: 15061111). Folic acid concentration was determined based on the standard curve generated using the folic acid standards.

### FFPE tissue block preparation

Male P21 pups were euthanized via CO2 inhalation. Mouse brain tissue were then carefully dissected, cut into 4mm thick blocks, and fixed in 10% neutral buffered formalin (NBF) at RT overnight. Subsequently, the tissue blocks were transferred into cassettes, dehydrated by a serial concentration of ethanol, and embedded in paraffin. These formalin-fixed paraffin-embedded (FFPE) tissue blocks were stored at 4°C until future use. To assess the quality of FFPE tissue blocks, RNA was extracted from ten 10 μm-thick FFPE sections using the Quick-RNA Mini Prep Plus Kit (Zymo Research, cat# R1057) following the manufacturer’s instructions. The quality of RNA was assessed by determining the percentage of total RNA fragments >200 nucleotides (DV200) using High-Sensitivity RNA ScreenTape on a 4150 TapeStation (Agilent). Tissue blocks with DV200 > 50% were considered to be of good quality and used for FFPE Visium spatial transcriptomics library preparation.

### FFPE Visium spatial transcriptomics library preparation

FFPE Visium Spatial transcriptomics libraries were prepared following the Visium Spatial Gene Expression Reagent Kits for FFPE User Guide (10x Genomics, cat# CG000407). The FFPE tissue blocks were sectioned at a thickness of 5μm using the HistoCore MultiCut microtome (Leica, cat# 149Multi0C1). These sections were spread on the water bath at 39°C and placed onto the Visium Spatial Gene Expression Slides. After drying on a thermo cycler at 42°C for 3h and at RT overnight in a desiccator, the tissue sections were further incubated on a thermo cycler at 60°C for 2h and deparaffinized by xylene and a serial of ethanol concentrations. Hematoxylin and eosin (H&E) staining was performed, and brightfield histology images were taken using an 80x objective on the MoticEasy Slide scanner. The tissue sections were then de-crosslinked, permeabilized, and hybridized with mouse WT probes. After hybridization at 50°C overnight and post-hybridization washes, probe ligation and post-ligation wash steps were conducted. Sequential processes of RNA digestion, probe release, probe extension, and probe elution were carried out. A fraction (1 μL) of the eluate was used for qPCR to determine the suitable PCR cycles, while the remaining elute (∼45μL) were utilized for sample index PCR. The PCR products were purified using SPRIselect beads. The quality of the libraries was assessed by Qubit 3.0 Fluorometer (Thermo Fisher, cat# Q33216) and the D100 DNA tapestation (Agilent). Finally, the libraries were pooled and sequenced on a NovaSeq 6000 System (Illumina) to obtain paired-end 150 bp reads.

### FFPE Visium spatial transcriptomics data analysis

The FFPE Visium sequencing reads were mapped to the mouse reference genome (mm10) using the short-read probe alignment algorithm for FFPE ‘count’ method in Space Ranger (v2.0.0) from the 10x Genomics. The resulting gene-count matrices and associated H&E images were then registered using the R package Seurat (v5.0.3) for subsequent analysis. The FFPE Visium spots with fewer than 500 UMIs or 250 genes were removed. The remaining gene-count matrices were normalized using the counts per million (CPM) method and transformed using the natural logarithm. These normalized gene-count matrices were then merged into a single object for joint processing and analysis.

To eliminate sample-related noise, the Harmony method was applied, and the harmonized data was clustered using the 3,000 most variable genes with the FindNeighbors function. Marker genes for each cluster were identified using the FindAllMarkers function, which performs a pairwise Wilcoxon Rank Sum test comparing the spots within each cluster against all other spots in the dataset. The resulting clusters were manually annotated based on both tissue histology and the presence of known marker genes for various brain regions/cell types.

Differential gene expression analysis between the MF and HF groups was conducted for each cluster using the non-parametric Wilcoxon Rank Sum test. Differentially expressed genes (DEGs) were identified based on a threshold of a 1.2-fold change and an adjusted p-value of 0.05.

### Single-nucleus multiome library preparation

Mouse hippocampal nuclei were isolated following the 10x Genomics protocol with slight modifications. P21 male mice were anesthetized and the hippocampi were dissected on a petri dish containing ice-cold PBS under a microscope. The dissected hippocampi were homogenized using a glass dounce homogenizer in NE buffer (0.32M sucrose, 10 mM Tris-HCl pH 8.0, 5 mM CaCl2, 3 mM MgCl2, 1 mM DTT, 0.1 mM EDTA, 0.1% NP40, 1x proteinase inhibitor cocktail, 1x RNase inhibitor). The homogenate was filtered through 70-μm cell strainers to remove tissue debris. After centrifugation, the pellet was resuspended in NE buffer, and 50% iodixanol solution (containing 20mM Tris-HCl pH 8.0, 25mM KCl, and 5mM MgCl2) was added to achieve a final iodixanol concentration of 30%. The mixture was centrifuged at 6,000g at 4°C for 20min. The cell debris on top was removed by aspiration, and the debris-free nuclei pellet at the bottom was gently resuspended in 100μL 0.1x lysis buffer and permeabilized on ice for 2min. The nuclei suspension was washed and centrifuged to obtain the nuclei pellet, which was then resuspended in diluted nuclei buffer supplemented with 1x RNase inhibitor. Approximately 6,000 nuclei per sample were loaded onto the 10x Genomics platform.

Single nucleus RNA-seq and single nucleus ATAC-seq libraries were simultaneously constructed using the Chromium Next GEM Single Cell Multiome ATAC + Gene Expression Kit (10x Genomics, cat# 1000285) according to the manufacturer’s instructions. The concentration of the libraries was measured by Qubit 3.0 Fluorometer (ThermoFisher, cat# Q33216) and the size distribution was assessed using D1000 screen tape on the 4150 tapestation system (Agilent, cat# G2992AA). The libraries were pooled for sequencing on the Illumina NovaSeq 6000 platform to obtain paired-end 150 bp reads.

### Single-nucleus multiome data analysis

The raw sequencing reads were trimmed and aligned to the mouse reference genome (mm10) using Cell Ranger-Arc (10x genomics, v.2.0.1) and quantified using cellranger-arc count function. The resulting count matrix was further processed using the R package Seurat (v.5.0.3) for snRNA-seq and Signac (v.1.13.0) for snATAC-seq. For quality control, the following filtering steps were applied: for snRNA-seq, nuclei with the number of genes (nFeature_RNA) ranging from 500 to 7000 and less than 2% mitochondrial genes were retained; for snATAC-seq, nuclei with unique reads ranging from 1000 to 70,000, a nucleosome signal < 4, and a TSS enrichment score > 2 were retained. Only nuclei passing quality control for both snRNA-seq and snATAC-seq were used for downstream analysis.

Based on the 3,000 most variable genes, the normalized snRNA-seq count matrix was used for PCA analysis. The first 20 PCs (Standard Deviation > 3) were used for uniform manifold approximation and projection (UMAP) and shared nearest neighbor (SNN) computation, identifying 14 clusters. For snATAC-seq normalization and linear dimensional reduction, the Signac functions FindTopFeatures, RunTFIDF, and RunSVD were utilized. Latent semantic indexing (LSI) dimensions 2-15 were used for anchoring strategy-based sample integration and UMAP clustering with 12 clusters identified. To integrate and simultaneously measure multiple modalities (RNA + ATAC), a weighted nearest neighbor (WNN) approach was employed. For this, 1-13 PCs from RNA and 2-12 LSIs from ATAC were used to construct a WNN graph, resulting in 13 clusters.

### Gene activity scoring, differential expression analysis and trajectory analysis

Genomic information for ATAC peaks was annotated using R package annotatr (v.1.28.0) [50] with the built-in mm10 genome. Additionally, ChIPseeker (v.1.38.0) [51] and TxDb.Mmusculus.UCSC.mm10.knownGene (v.3.10.0) [52] were used for further genomic annotation. Gene activity scores for each cell were computed from snATAC-seq data by calculating an exponential decay weighted sum of fragment counts from each gene’s transcription start site (TSS) to transcription terminal site (TTS) using Signac (v.1.13.0) [53]. Raw gene scores were normalized by dividing each score by the mean gene activity score per cell. To identify differentially expressed genes (DEGs), the FindMarker function in R package Seurat (v.5.0.3) [37] was utilized with Wilcoxon Rank Sum test. DEGs were identified with thresholds set at a 1.2-fold change and an adjusted p-value of 0.05.

Trajectory analysis was performed to understand the dynamic progression of cells along a developmental path of DG neurons. Monocle3 (v.1.3.1) [54] was utilized for trajectory and pseudotime analysis. The subset of DG neuron Seurat object was transformed and passed to the Monocle object. Dimensionality reduction was performed using Monocle3 independent of the result from Seurat. The trajectory was inferred by applying the Monocle3::learn_graph function and cells were ordered along the trajectory using the Monocle3::order_cells function. Monocle3::graph_test was used to identify pseudotime correlated genes, adjusted p-value < 0.05.

### RT-qPCR

Total RNA samples were extracted from P21 male hippocampus tissues using TRIzol/chloroform phase separation combined with column purification using RNA clean and concentrator-25 kit (Zymo Research, cat#). DNase I digestion on columns was performed to eliminate residual DNA contamination. 1 μg total RNA was reverse transcribed to cDNA using High-Capacity cDNA Reverse Transcription Kit (Thermo Fisher, cat# 4368814). Realtime qPCR was performed using GoTaq™ qPCR Master Mix (Promega, cat# A6001) targeting the following mouse genes: Rbfox3 (forward: CACTCTCTTGTCCGTTTGCTTC, reverse: CTGCTGGCTGAGCATATCTGTA), Gfap (forward: ACCAGTAACATGCAAGAGACAGAG, reverse: GATAGTCGTTAGCTTCGTGCTTG), Olig2 (forward: GGTGGTACCGGTGCAGCAACTGCCACTAAGTA, reverse: GGTGGTGTCGACTCTGGACCGGAGATCTGAATAG) and GAPDH (forward: AATGGTGAAGGTCGGTGTG, reverse: GTGGAGTCATACTGGAACATGTAG).

## Code availability

https://github.com/YU-LIN96/Maternal_HF_brain_development

## Data availability

The Visium spatial transcriptomics datasets were deposit at NCBI with accession number GSE269543. The single-nucleus multiome (snRNA-seq and snATAC-seq) datasets were deposit at NCBI with access number GSE269544.

## Ethics approval and consent to participate

All procedures were performed following national and international guidelines and approved by the institutional board at Virginia Tech. The study is in accordance with National Institute of Health guidelines.

## Author contribution statement

H. X. conceived the experimental design. X. X., P. da. S. S., B. C., C. P., F. C., T. H., S. F., and K. Z. performed the experiments. X. X., Y. L., and L. Y. analyzed the data. X. X., Y. L., L. Y., X. W., and H X interpreted the results and drafted the manuscript. All authors discussed the results, read, and edited the manuscript, and approved the final manuscript.

---

**Supplementary Figure 1.**
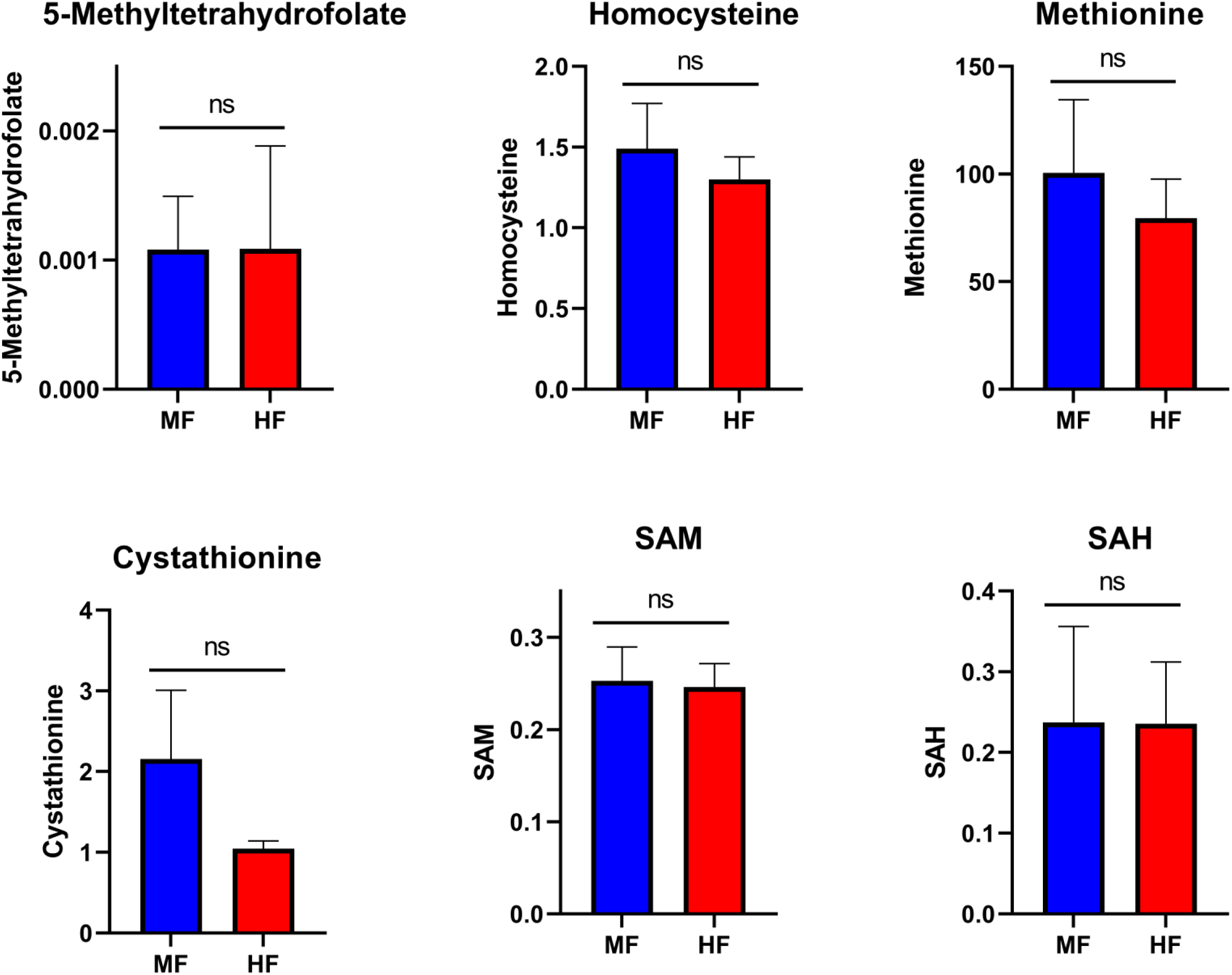
Quantification of one-carbon metabolites by LC-MS. Maternal plasma samples were collected when the pups reached P21 and analyzed using LC-MS to quantify one-carbon metabolites: Methyltetrahydrofolate (5-Me-THF), homocysteine (Hcy), methionine (MET), cystathionine (CYSTA), S-adenosylmethionine (SAM), and S-adenosyl-L-homocysteine (SAH). ns. not significant.

**Supplementary Figure 2.**
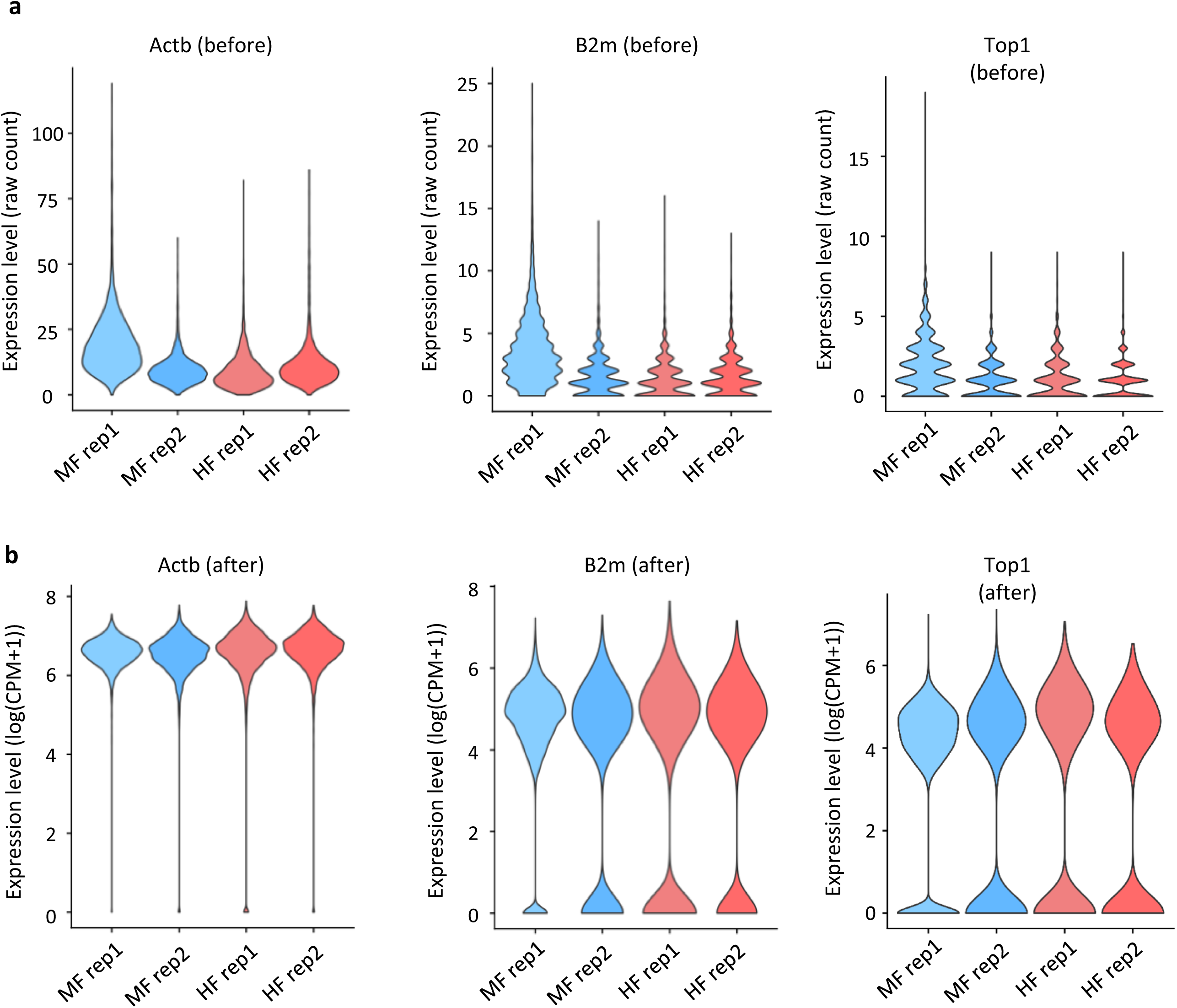
Normalization of Visium spatial transcriptomics datasets. **a**. Violin plot showing the raw counts of the house keeping genes Actb, B2m and Top1 before data normalization. **b**. Violin plot showing the normalized and log transformed counts of Actb, B2m and Top1.

**Supplementary Figure 3.**
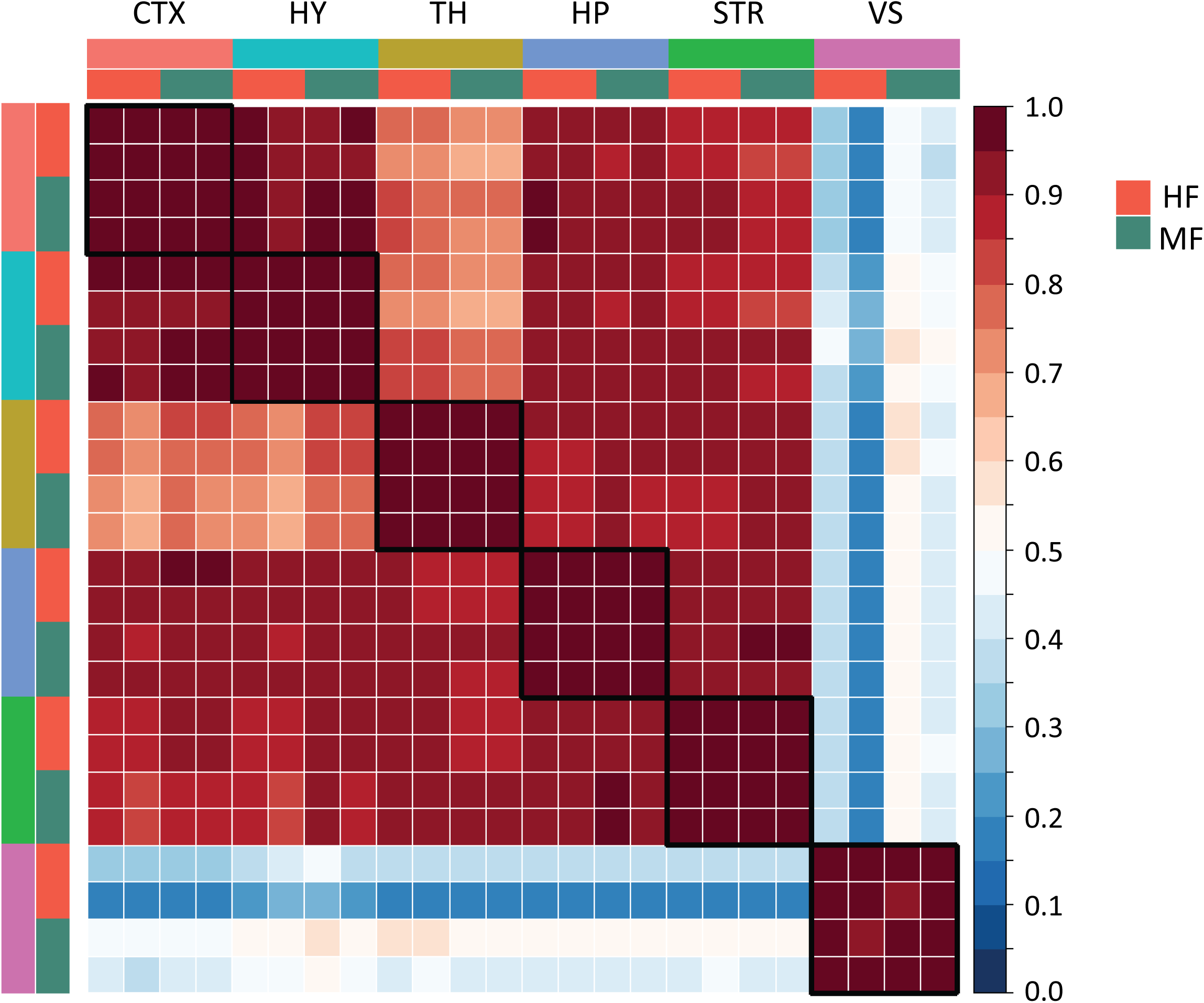
Replicate reproducibility. Heatmap showing the correlation of spatial transcriptomic profiles between two biological replicates across six brain regions for both MF and HF groups.

**Supplementary Figure 4.**
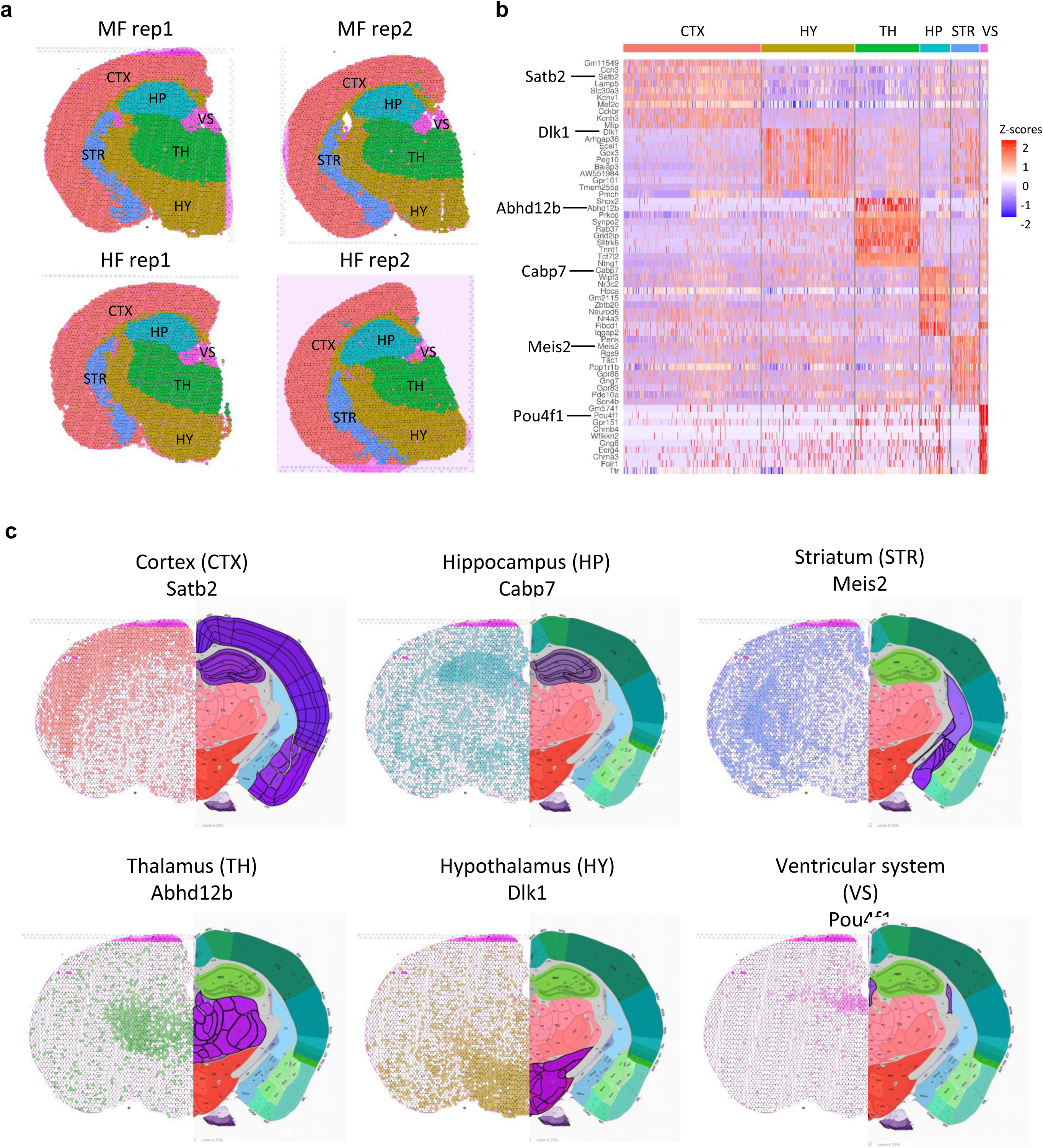
FFPE Visium spatial transcriptomics agreement with anatomical regions. **a**. Annotation of the six major clusters based on their co-localization with anatomical landmarks from the mouse P56 coronal section #72 of the Allen Mouse Brain Atlas. **b**. Heatmap showing the top 10 marker genes for each brain region. **c**. Spatial density plot of representative marker genes for each brain region (left) and the corresponding anatomical landmarks (right).

**Supplementary Figure 5.**
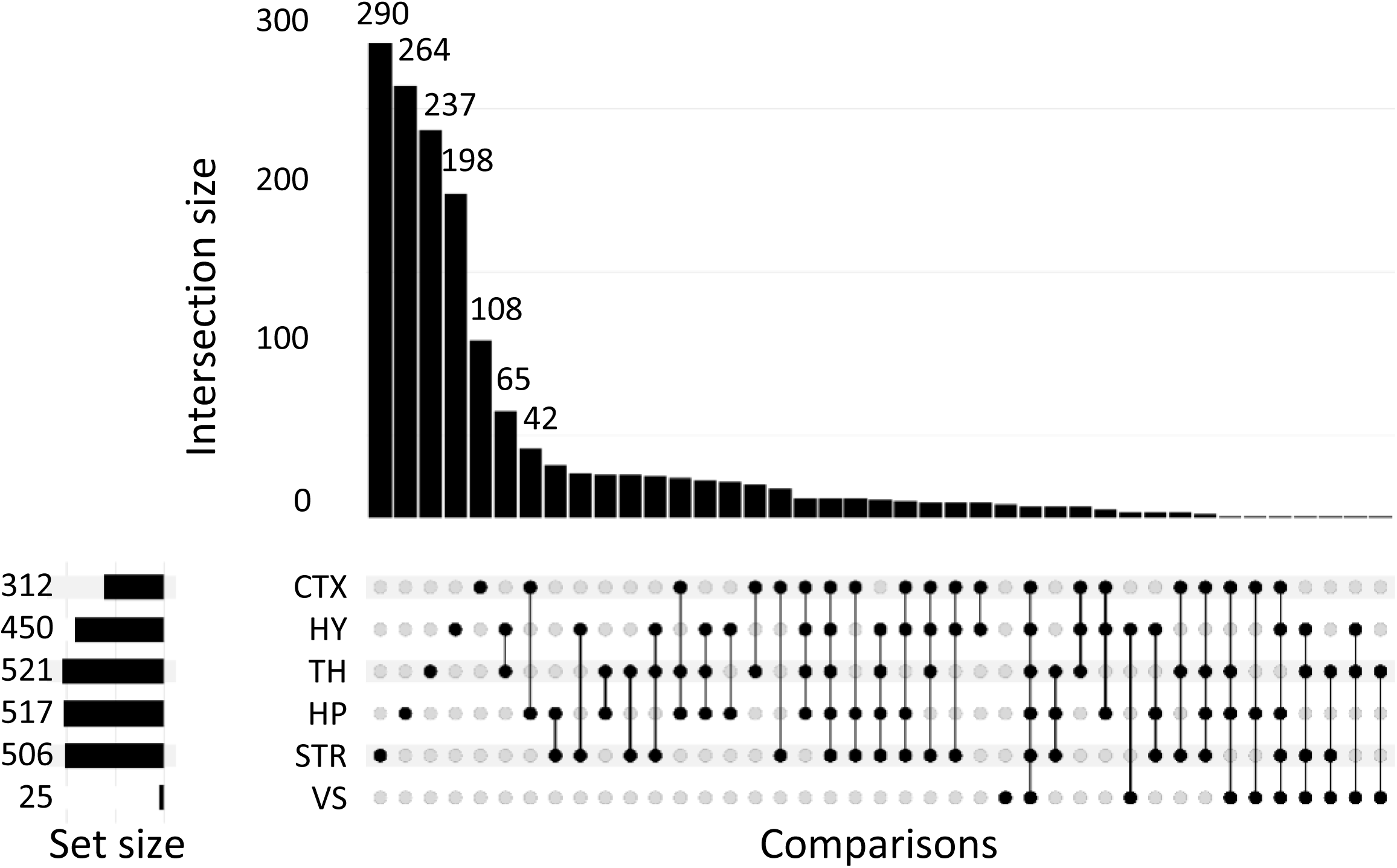
Shared and region-specific DEGs among various brain regions. Intersection bar plot showing the number of shared and brain region-specific differentially expressed genes. The horizontal bar on the bottom left side shows the number of DEGs. Different intersection combinations of DEGs are represented by the dot plot. The vertical bar plot shows the number DEGs in the indicated combination of intersection.

**Supplementary Figure 6.**
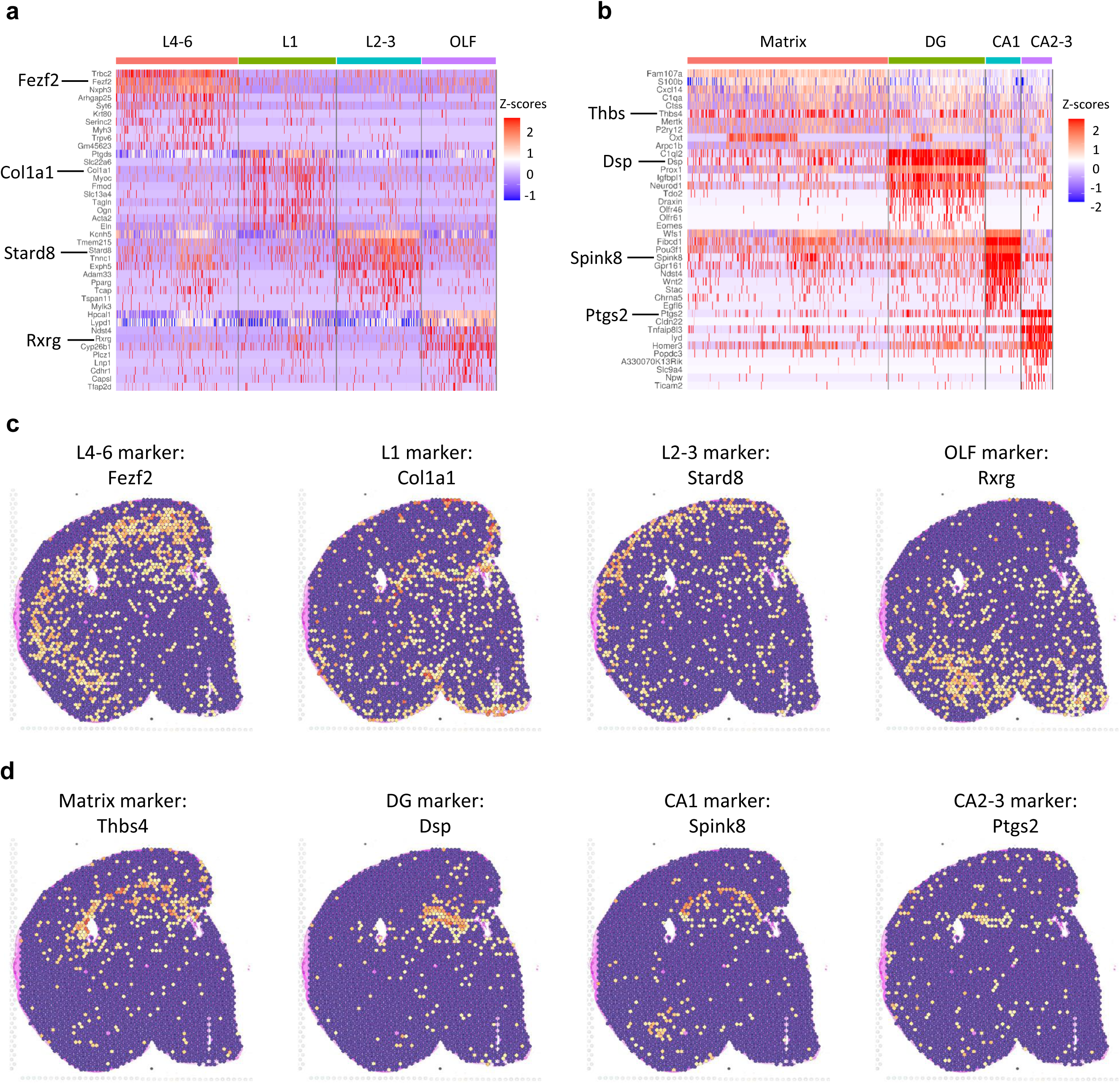
FFPE Visium spatial transcriptomics aligns subregions in the cortex and hippocampus. **a**. Heatmap displaying the top 10 marker genes for each sub-cluster in the cortex region. **b**. Heatmap showing the top 10 marker genes for each sub-cluster in the hippocampus region. **c**. Spatial density plot of representative marker genes for each sub-cluster in the cortex region. **d**. Spatial density plot of representative marker genes for each sub-cluster in the hippocampus region.

**Supplementary Figure 7.**
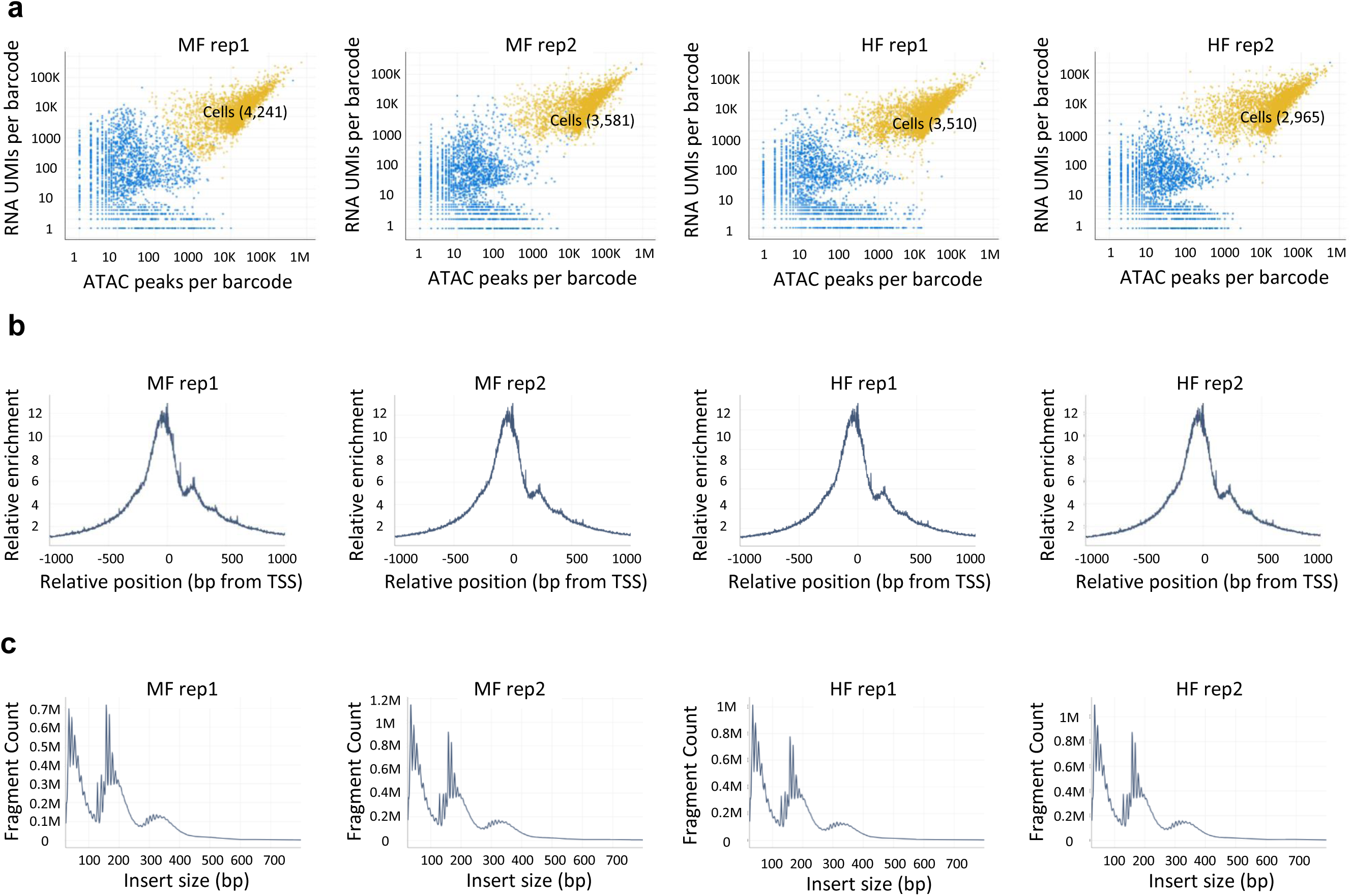
Confirmation of the quality of single-nucleus multiome libraries. **a**. Scatter plot showing the joint cell calling using the cross sensitivities between ATAC transposition events in peaks per barcode and RNA UMIs per barcode. Spots representing cells were highlighted in yellow and the number of cells for each sample was labelled. **b**. Enrichment of ATAC signal around the transcription start sites (TSS) in the four samples. **c**. Distribution of fragment size for nuclei in each sample that passed the quality control threshold.

**Supplementary Figure 8.**
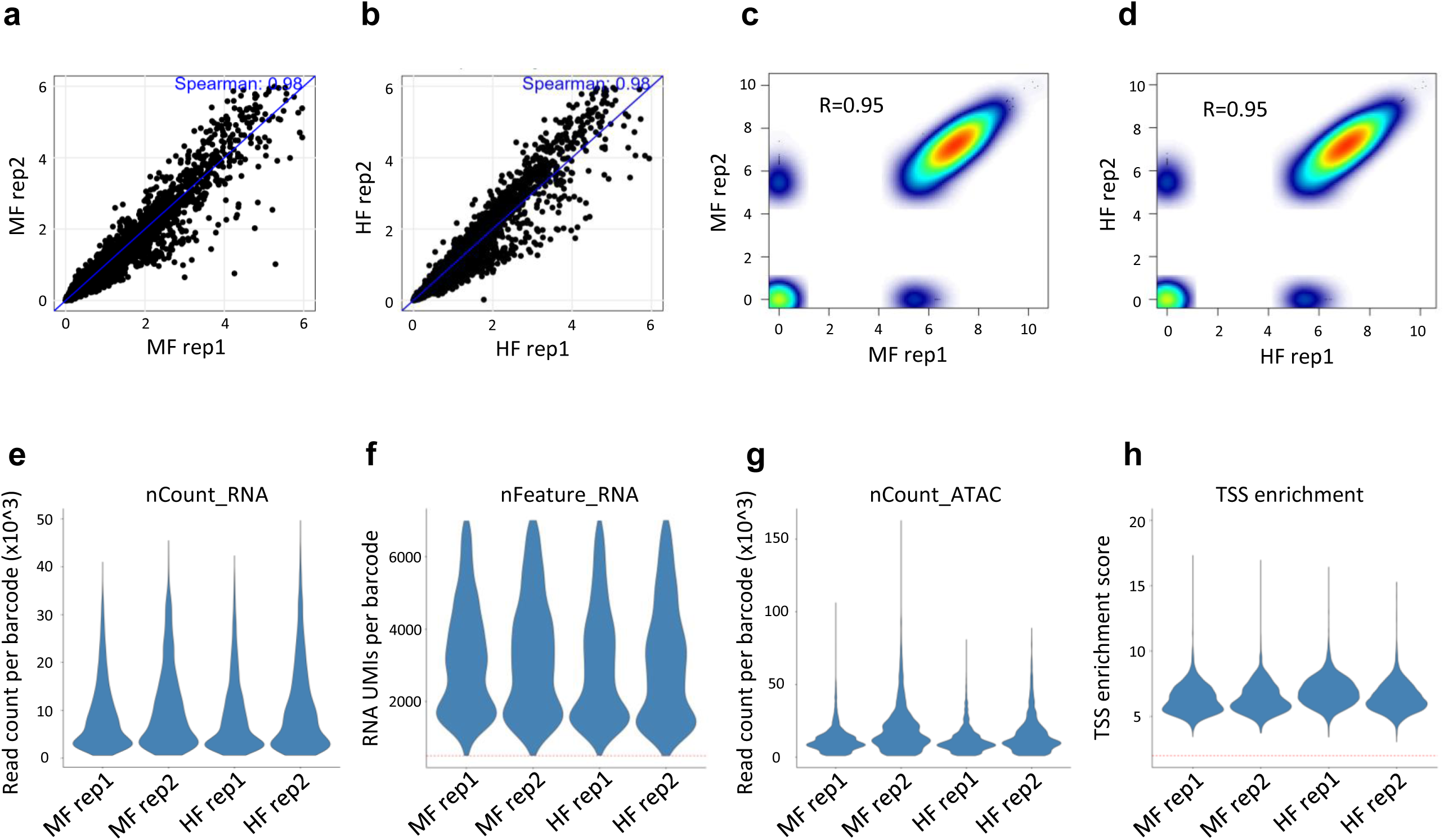
Quality control of single-nucleus multiome datasets. **a,b**. Dot plot showing the correlation of snRNA-seq datasets between two biological replicates, represented by log2(normalized counts) in the MF (**a**) and HF (**b**) groups. **c, d**. Dot plot showing the correlation of snATAC-seq datasets between two biological replicates, represented by log2(RPKM), in the MF (**c**) and HF (**d**) groups. **e**. Violin plot showing the nCount (number of reads) of snRNA-seq datasets. **f**. Violin plot showing the nFeature (number of genes) of snRNA-seq datasets. **g**. Violin plot showing the nCount (number of reads) of snATAC-seq datasets. **h**. Violin plot showing the TSS enrichment scores of snATAC-seq datasets.

**Supplementary Figure 9.**
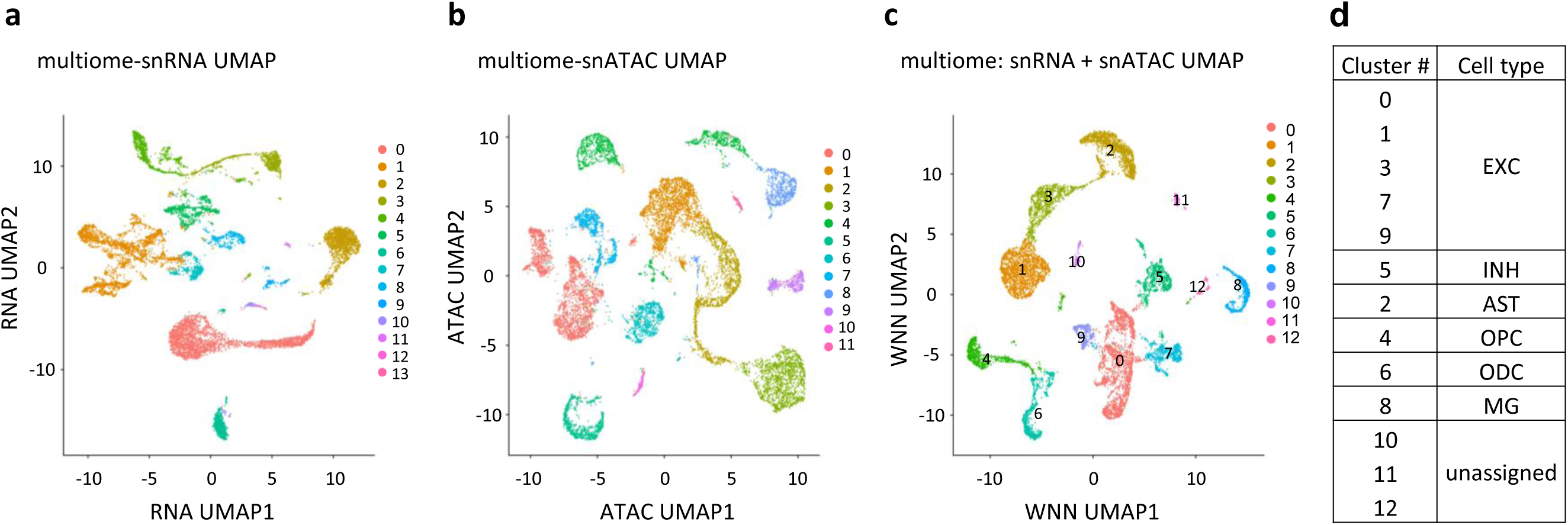
Clustering of single-nucleus multiome datasets. **a**. UMAP visualization of snRNA-seq datasets. **b**. UMAP visualization of snATAC-seq datasets. **c**. UMAP visualization of combined snRNA-seq and snATAC-seq datasets using the WNN approach. Nuclei are colored based on clusters identified in each clustering strategy. **d**. Clusters identified in the WNN approach were assigned to major cell types.

**Supplementary Figure 10.**
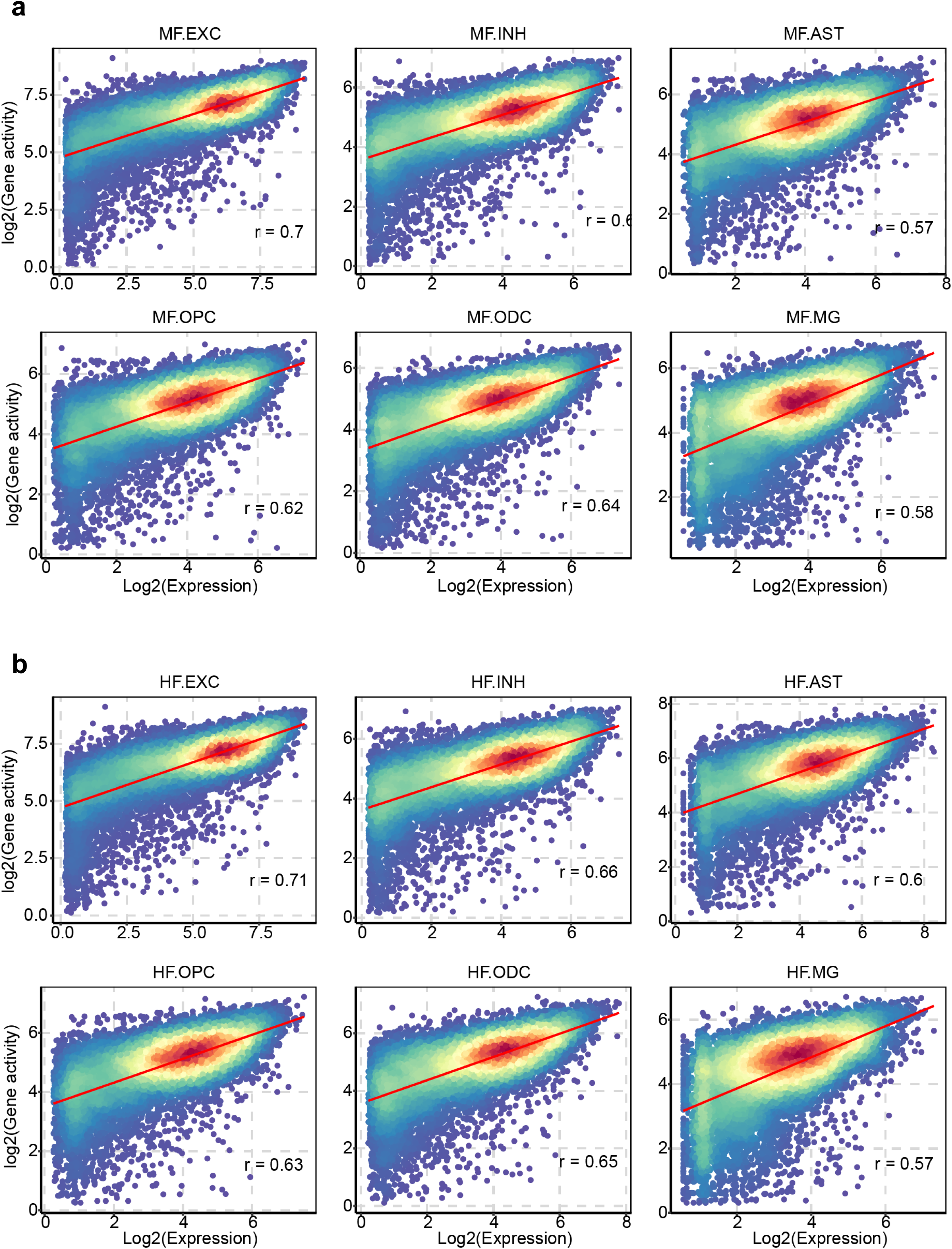
Correlation between gene expression and gene accessibility in major cell types. **a**. Correlation between gene expression and gene accessibility for each cell type in the MF group. **b**. Correlation between gene expression and gene accessibility for each cell type in the HF group.

**Supplementary Figure 11.**
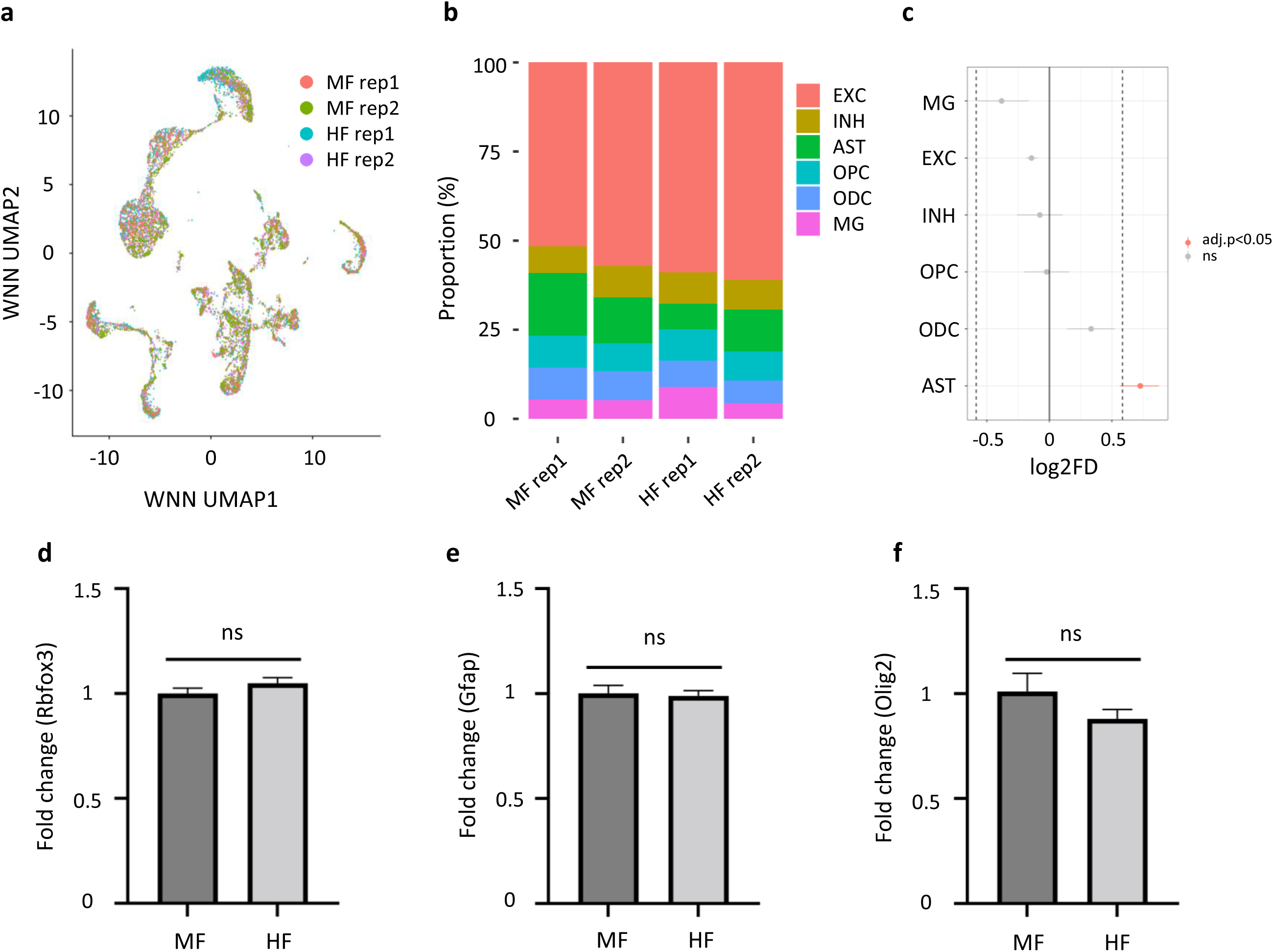
Proportion of major cell types identified with the single-nucleus multiome. **a**. Integrated UMAP visualization of snRNA-seq and snATAC-seq datasets colored by samples analyzed. **b**. Bar plot showing the relative percentage of each major cell type in each sample. **c**. Population shift represented by log2 fold change. **d, e, f**. Relative mRNA expression level of Rbfox3 (pan neuronal marker), Gfap (astrocyte marker), and Olig2 (Oligodendrocyte marker) in the hippocampus tissues of P21 male pups. Four biological replicates were included. Ns: not significant by t-test.

**Supplementary Figure 12.**
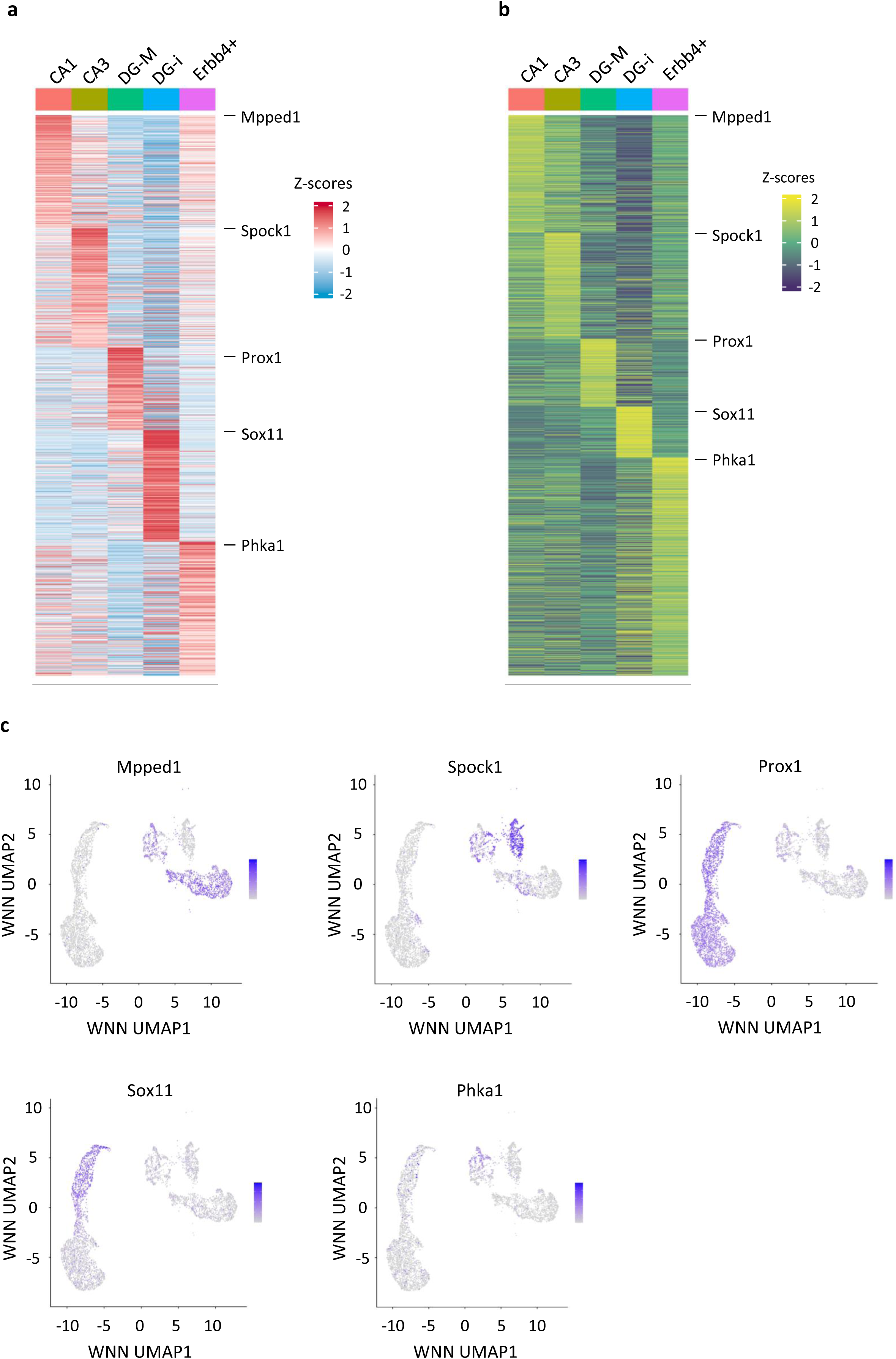
Marker genes identified in hippocampal excitatory neuron subtypes. **a, b**. Row-normalized heatmaps for single nucleus gene expression (a) or gene activity (b) of marker genes specific to excitatory neuron subtypes. **c**. UMAP density plot illustrating the signature genes for each excitatory neuron subtype.

**Supplementary Figure 13.**
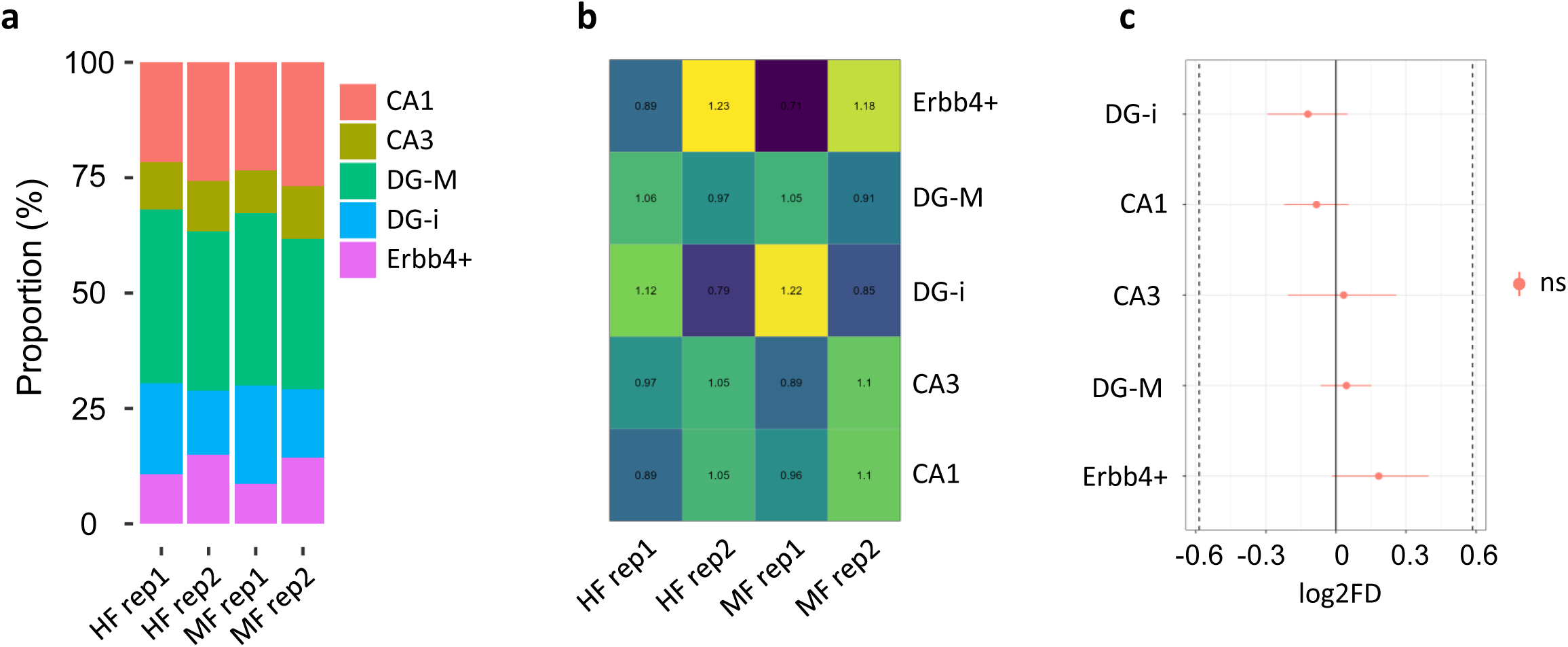
Proportion of hippocampal excitatory neuron subtypes identified from the single-nucleus multiome datasets. **a**. Bar plot showing the relative percentage of each excitatory neuron subtype in each sample. **b**. Heatmap showing the ratio of observed vs expected of each excitatory neuron subtype. **c**. Population shift represented by log2 fold change. ns: not significant.

**Supplementary Figure 14.**
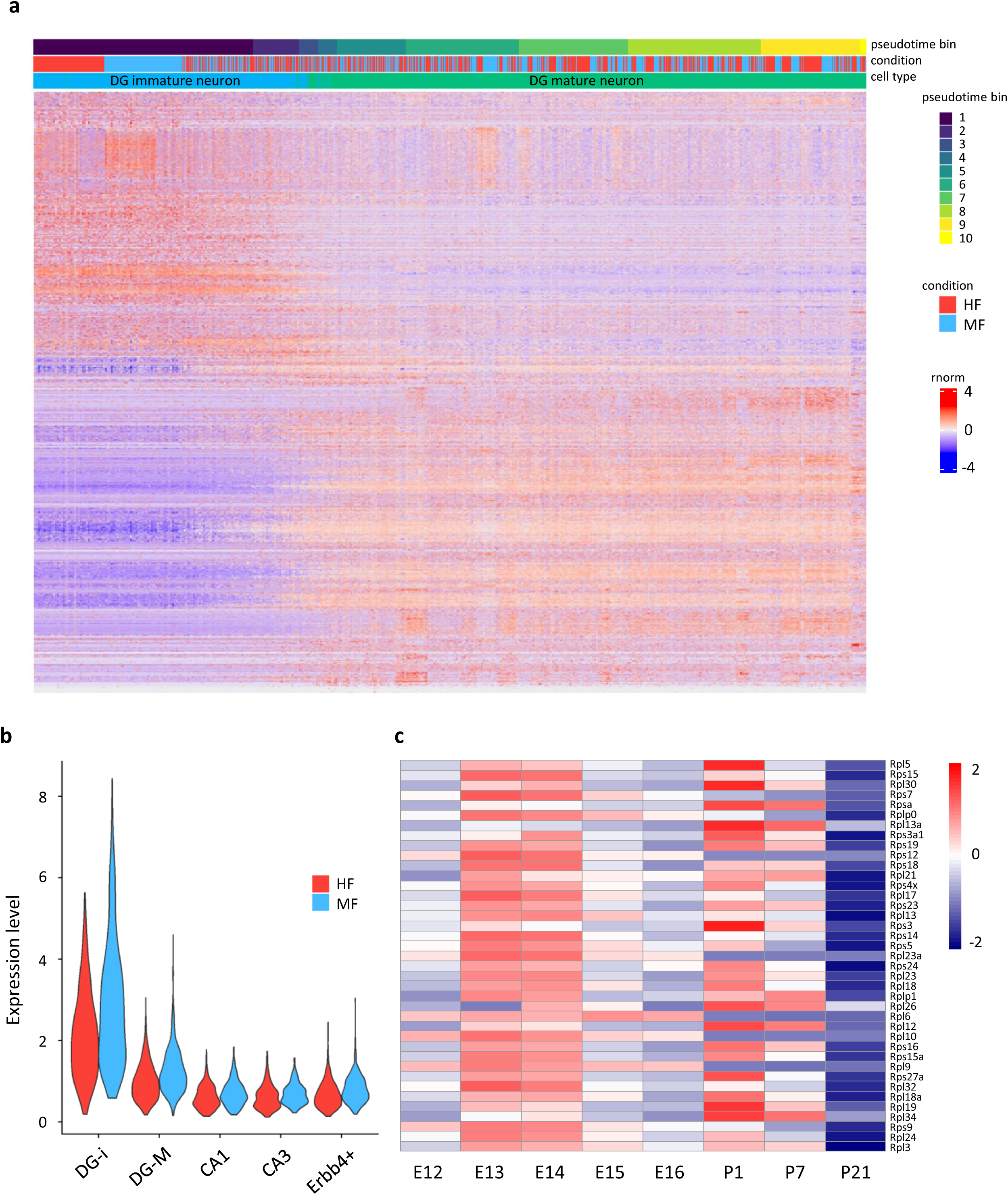
Expression profiles of ribosomal protein-coding genes in excitatory neurons. **a**. Heatmap showing the expression of genes along the trajectory of DG neurons. **b**. Violin plot showing the expression levels of ribosomal protein-coding genes that were both DEGs and trajectory-related in the MF and HF groups across hippocampal excitatory neuron subtypes. **c**. Heatmap showing the expression of trajectory-related ribosomal protein-coding DEGs in excitatory neurons during embryonic and postnatal brain development.

**Supplementary Figure 15.**
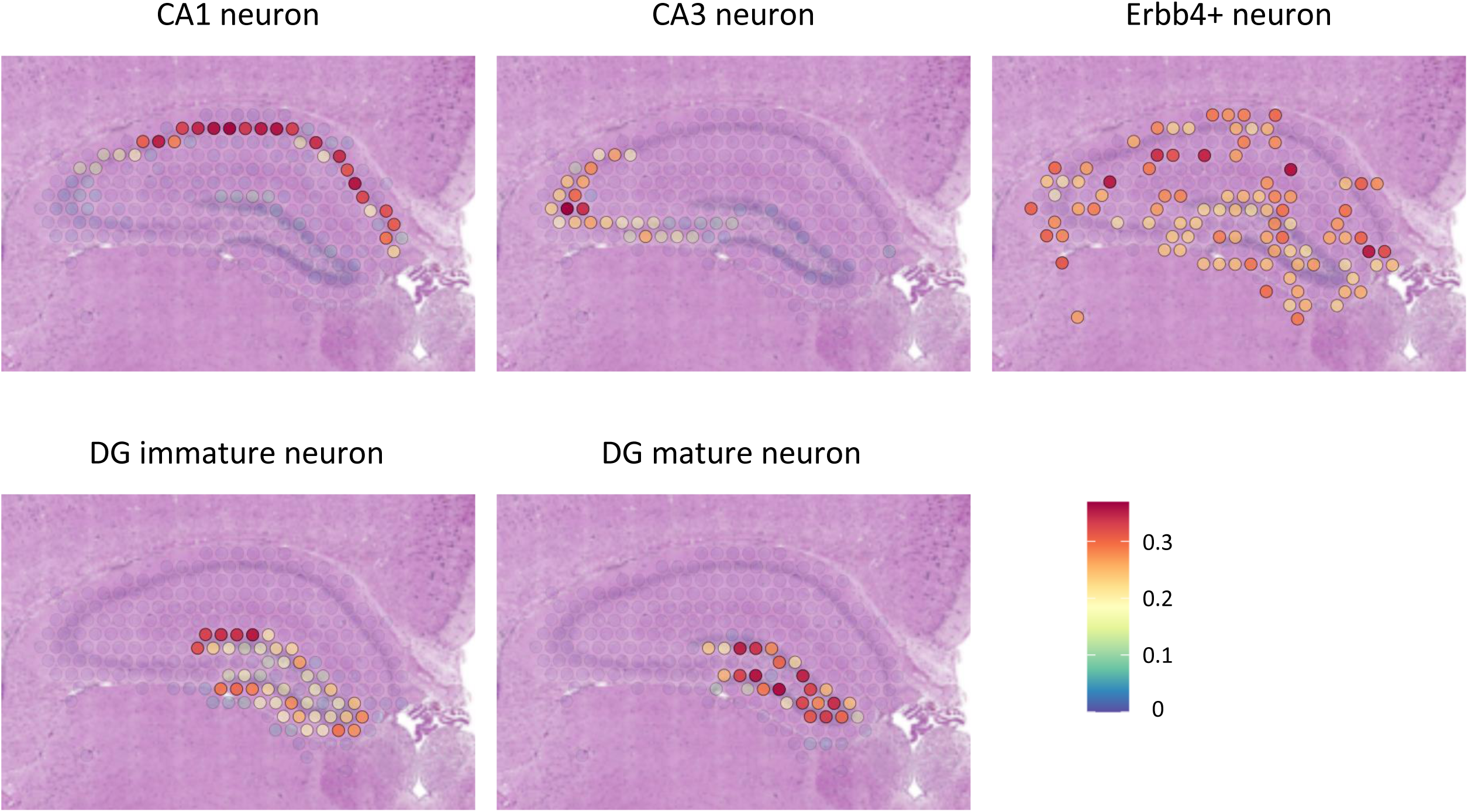
Projection of snRNA-seq to spatial transcriptomics. Projection of the hippocampal excitatory neuron subtypes to spatial transcriptomics: CA1 neurons, CA3 neurons, Erbb4+ neurons, DG immature neurons, and DG mature neurons.

**Supplementary Table 1. Mapping statistics of spatial transcriptomics and single-nucleus multiome libraries.**

**Supplementary Table 2. Differentially expressed genes identified in major brain regions.**

**Supplementary Table 3. Differentially expressed genes identified in cortex and hippocampal subregions.**

**Supplementary Table 4. Differentially expressed genes identified in major hippocampal cell types.**

**Supplementary Table 5. Differentially expressed genes identified in hippocampal excitatory neuron subtypes.**

